# Expansion of Functional Regulatory T Cells Using Soluble RAGE Prevents Type 1 Diabetes

**DOI:** 10.1101/2020.01.10.902627

**Authors:** Sherman S. Leung, Danielle J. Borg, Domenica A. McCarthy, Tamar E. Boursalian, Justen Cracraft, Aowen Zhuang, Amelia K. Fotheringham, Nicole Flemming, Thomas Watkins, John J. Miles, Per-Henrik Groop, Jean L. Scheijen, Casper G. Schalkwijk, Raymond J. Steptoe, Kristen J. Radford, Mikael Knip, Josephine M. Forbes

## Abstract

Type 1 diabetes (T1D) is an autoimmune disease with no cure. Therapeutic translation has been hampered by preclinical reproducibility. Here, short-term administration of an antagonist to the receptor for advanced glycation end products (sRAGE) protected against murine diabetes at two independent centers. Treatment with sRAGE increased regulatory T cells (T_regs_) within islets, pancreatic lymph nodes and spleen, increasing islet insulin expression and function. Diabetes protection was abrogated by T_reg_ depletion and shown to be dependent on antagonizing RAGE using knockout mice. Human T_regs_ treated with a RAGE ligand downregulated genes for suppression, migration and T_reg_ homeostasis (*FOXP3, IL7R, TIGIT, JAK1, STAT3, STAT5b, CCR4*). Loss of suppressive function was reversed by sRAGE, where T_regs_ increased proliferation and suppressed conventional T cell division, confirming that sRAGE expands functional human T_regs_. These results highlight sRAGE as an attractive treatment to prevent diabetes, showing efficacy at multiple research centers and in human T cells.

## Introduction

Type 1 diabetes (T1D) is an autoimmune disease involving a heterogeneous interplay between genetics and environmental factors resulting in T cell mediated destruction of insulin-producing β cells (Atkinson et al., 2014). T1D incidence is increasing at 2-3% per year worldwide, elevating the risk for premature death and costing $14 billion per annum in healthcare in the USA alone (Tao et al., 2010). Risk factors for developing T1D include decreases in functional regulatory T cells (T_regs_) (Brusko et al., 2005) and increased numbers of autoantigen-specific conventional T cells (T_convs_), particularly those with an effector phenotype (T_eff_) (Laban et al., 2018). Early phase clinical trials promoting the expansion of T_regs_, thereby suppressing T_conv_ activation, have shown promise in preserving insulin production after the diagnosis of T1D (Alhadj Ali et al., 2017; Rosenzwajg et al., 2015), but await validation in larger cohorts. However, it is now recognized that interventions aimed at reversing clinically diagnosed T1D may be “too late”, with Phase III clinical trials not reaching primary end points (Insel et al., 2015). As a result, promising therapeutics targeting T cells are being repurposed for use prediabetes to prevent the onset of T1D (NCT01030861; NCT01773707). Recently, the former study, which was a phase II randomized trial, reported that Teplizumab – a Fc-receptor non-binding anti-CD3 monoclonal antibody – significantly delayed the onset of T1D (Herold et al., 2019).

In fact, a framework for staging prediabetes was recently established which involves standardized screening to accurately stratify individuals into Prediabetes Stages 1-3 so that novel therapies can be tested within a suitable therapeutic window (Insel et al., 2015). Thus, interventions delivered prediabetes are clinically feasible, have a greater chance of preserving insulin secretion and could prevent the onset of T1D.

The receptor for advanced glycation end products (RAGE) is a pattern recognition receptor implicated in inflammatory disease and is expressed in various cells involved in T1D including T cells (reviewed in (Leung et al., 2016)). Recently, changes in RAGE expression have been associated with the risk for developing T1D in humans (Salonen et al., 2015; Salonen et al., 2014; Salonen et al., 2016). Furthermore, T cells from at-risk individuals who progress to T1D, have greater RAGE expression, which enhances T cell cytokine production and survival (Durning et al., 2016). Natural history studies have further revealed that polymorphisms in the RAGE gene (*AGER*) decrease circulating soluble RAGE (sRAGE) concentrations (Salonen et al., 2014), which is a naturally occurring antagonist that competes for RAGE ligands, increasing the risk of T1D (Forbes et al., 2011). The decreases in circulating sRAGE also coincide with seroconversion to autoantibodies against islet auto-antigens in at-risk individuals (Salonen et al., 2015; Salonen et al., 2016). Therefore, this deficiency in circulating sRAGE is a novel therapeutic target for preventing the onset of T1D.

Here, we targeted the deficiency in circulating sRAGE levels prediabetes by short-term administration of recombinant human sRAGE with the aim of preventing diabetes onset in mice. We show that it acts in an immunomodulatory manner to decrease diabetes incidence at two independent research centers through T_reg_ modulation. Additionally, sRAGE increased the proportion of T_regs_ in the islet infiltrating leukocytes, pancreatic lymph nodes (PLN) and spleen, which reduced islet infiltration and preserved islet numbers, insulin expression and β cell function. *Ex vivo,* sRAGE promoted the expansion of human T_regs_ and reduced T_conv_ proliferation in co-culture, whereas T_regs_ cultured in the presence of the RAGE ligand, AGEs, had reduced suppressive function. Our data suggest that short-term delivery of sRAGE prediabetes is an effective modulator of functional T_reg_ expansion and has efficacy to prevent diabetes and possibly other autoimmune diseases.

## Results

### Short-Term sRAGE Treatment Provides Lasting Protection from Diabetes in Multi-Site Trial

There is no cure for T1D so there is a desperate need for reproducible and translatable novel disease targets. Here, we administered recombinant human sRAGE prediabetes to correct the deficiency in circulating sRAGE seen in at-risk progressors who develop T1D (Salonen et al., 2015; Salonen et al., 2014; Salonen et al., 2016), using a preclinical study design tested for reproducibility at two independent research centers (Figure 1A).

**Figure 1.**
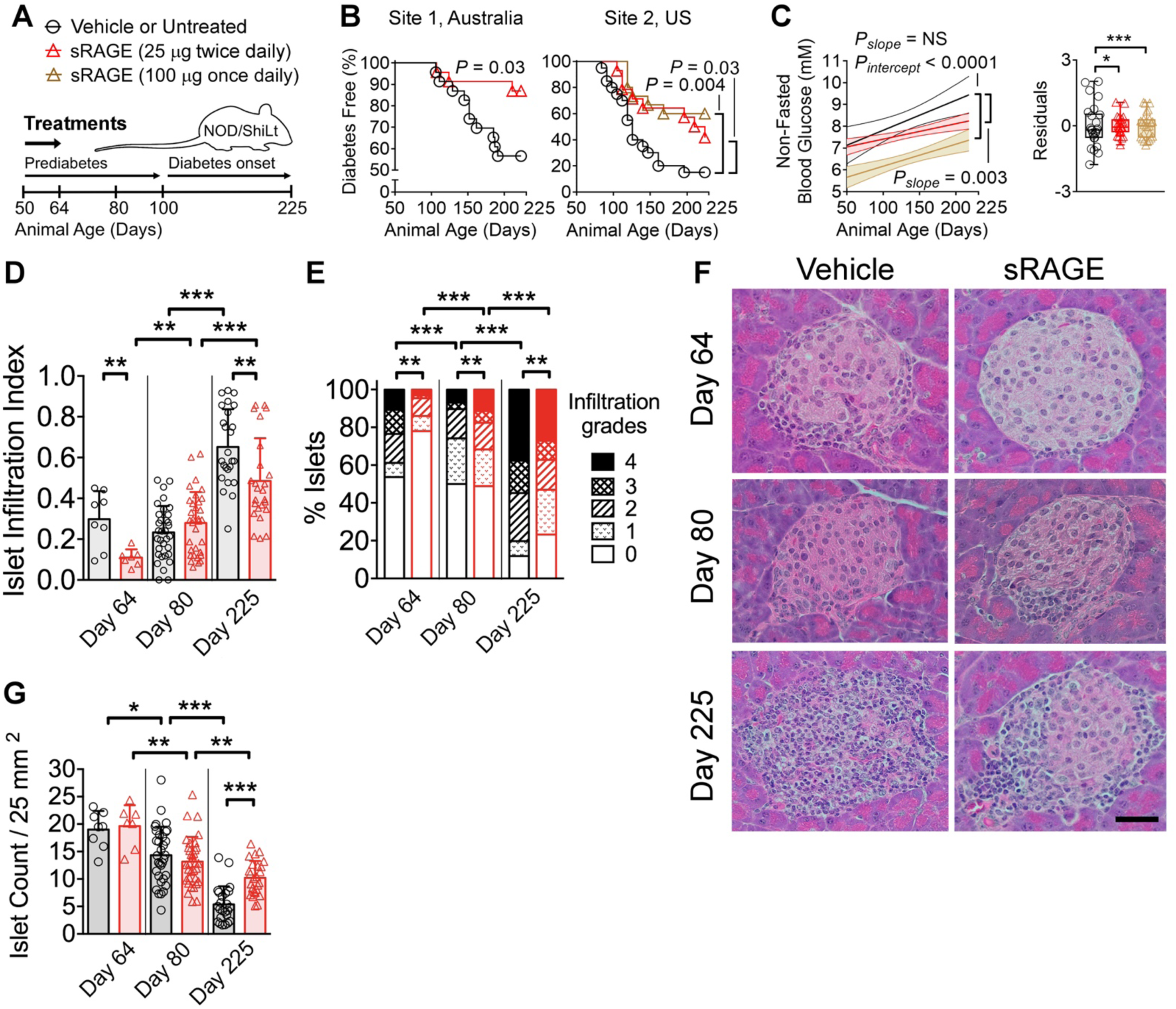
Treatment with sRAGE provides lasting protection against autoimmune diabetes in an international multi-site preclinical trial. (A) NOD/ShiLt mice were administered vehicle at Site 1 or untreated at Site 2 (black bars/circles), or treated with 25 μg sRAGE twice daily (red bars/triangles) or 100 μg sRAGE once daily (brown bars/triangles) from days 50-64 of life. (B) Autoimmune diabetes incidence. Site 1 (3 independent experiments, *n* = 23/group). Site 2 (1 independent experiment, *n* = 14-20/group). (C) Non-fasted blood glucose concentrations shown as linear regression ± 95% confidence intervals (left) and residuals representing variability of blood glucose levels from the regression line (right). (D-F) Pancreatic islet infiltration. (D) Islet infiltration index (0 indicates no infiltration; to 1 indicates >75% infiltration). (E) Degree of islet infiltration (grade 0, none; grade 1, peri-infiltration; grade 2, <25% infiltration; grade 3, 25-75% infiltration; grade 4, >75% infiltration). (F) Representative H&E photomicrographs (*n* = 7-33 sections/group from *n* = 4-7 mice/group, scale bar = 40 µm). (G) Islet count normalized by tissue area. Column graphs are shown as median (IQR) and analyzed by two-tailed Mann-Whitney U-test. Box and whisker plot variances were analyzed by F-test. Degree of insulitis is shown as mean and proportions analyzed by Fischer’s exact test. Diabetes incidence are shown as Kaplan-Meier survival curves and were analyzed by log-rank test. \**P* < 0.05, \*\**P* < 0.01, \*\*\**P* < 0.001. sRAGE, soluble receptor for advanced glycation end products.

When sRAGE was administered intraperitoneally twice daily, the incidence of diabetes decreased 3-fold when compared with vehicle treated mice (Figure 1B, Site 1). We found comparable results at an independent research center with greater diabetes penetrance (Figure 1B, Site 2), where sRAGE treatment resulted in a 2.8-fold reduction in diabetes incidence (vs. untreated; Figure 1B). At a greater dose administered only once daily, sRAGE treatment was even more effective and decreased diabetes incidence by 4-fold (vs. untreated; Figure 1B).

Non-fasted blood glucose levels in sRAGE treated mice were significantly lower over the study duration when compared with vehicle and untreated mice using regression analysis (Figure 1C). Furthermore, the regression line and its confidence intervals for the once daily sRAGE treatment regimen were lower than those of the remaining groups, until approximately day 200 when both sRAGE groups had overlap (Figure 1C). Blood glucose variability was also reduced following sRAGE administration, as determined by decreased residuals from the regression line (vs. vehicle/untreated; Figure 1C). These findings demonstrate that short-term sRAGE treatment significantly reduces diabetes incidence and improves long-term blood glucose control. From herein, we characterize the effects of sRAGE given twice daily because we were able to observe significant protection from diabetes at this lower dose.

### sRAGE Therapy Decreases Islet Infiltration and Increases Islet Numbers

Immune cell infiltration into the pancreatic islets and islet destruction are pathological hallmarks of T1D (Atkinson et al., 2014), so we hypothesized that these facets of disease pathogenesis would be improved by sRAGE treatment. To this end, immediately after therapy completion on day 64, mice treated with sRAGE had a considerably reduced islet infiltration index as compared with vehicle (Figure 1D). This was due to a decrease in the numbers of islets presenting with high grade insulitis (>75% infiltrate, grade 4) and an increase in islets without insulitis (0% infiltrate, grade 0; Figure 1E and 1F). Interestingly, by day 80, the islet infiltration index did not differ between groups (Figure 1D) but unexpectedly, sRAGE treated mice had a small increase in the proportion of islets with grade 4 insulitis (>75% infiltration; Figure 1E and 1F). When sRAGE treated mice were examined on day 225, we found a significant reduction in the islet infiltration index (Figure 1D), an increased proportion of islets without immune cell infiltration and a reduced proportion of islets that scored 4 (vs. vehicle; Figure 1E and 1F). Islet numbers decreased over the study duration in both cohorts (Figure 1G), but sRAGE treatment preserved a greater number of islets by day 225 (Figure 1G). Islet numbers did not differ between groups at the other time points (Figure 1G).

### sRAGE Rapidly Increases T_reg_/T_eff_ Ratios in the PLN and Spleen

T_regs_ in the PLN and spleen regulate islet infiltration, influencing the development of diabetes (McNally et al., 2011). Thus, we tested whether sRAGE could influence the numbers of T_regs_ and T_convs_ in these lymphoid tissues, in addition to T_reg_/T_eff_ ratios (gating strategies in Figure S1). Immediately after sRAGE therapy on day 64, higher numbers of CD4^+^CD8^-^CD25^+^Foxp3^+^ T_regs_, as well as FoxP3^-^CD4^+^ and FoxP3^-^CD8^+^ T_convs_ were observed in the PLN and spleen (vs. vehicle; Figure S2A-S2C). This is consistent with previous reports showing that sRAGE treatment is immunomodulatory, since it can increase the numbers of monocytes, macrophages and B cells (Pullerits et al., 2006). However, there are no studies showing that sRAGE can expand the numbers of T cells, nor its specific effects on the T cell subsets of T_regs,_ T_convs_ and T_effs_.

Here, we found a superior increase in PLN T_reg_ numbers, resulting in elevated T_reg_/T_eff_ cell ratios in sRAGE treated mice (Figure 2A-2C), which suggested the local environment was more regulated. Despite the increase in splenic T_regs_, its ratio to T_eff_ cells remained unchanged on day 64 (Figure 2A-2C). There was no change in the activation status of FoxP3^-^CD4^+^ and FoxP3^-^ CD8^+^ T_convs_ following sRAGE treatment in either lymphoid compartments, as determined by the expression of the CD62L and CD44 adhesion molecules (Figure S2B and S2C).

**Figure 2.**
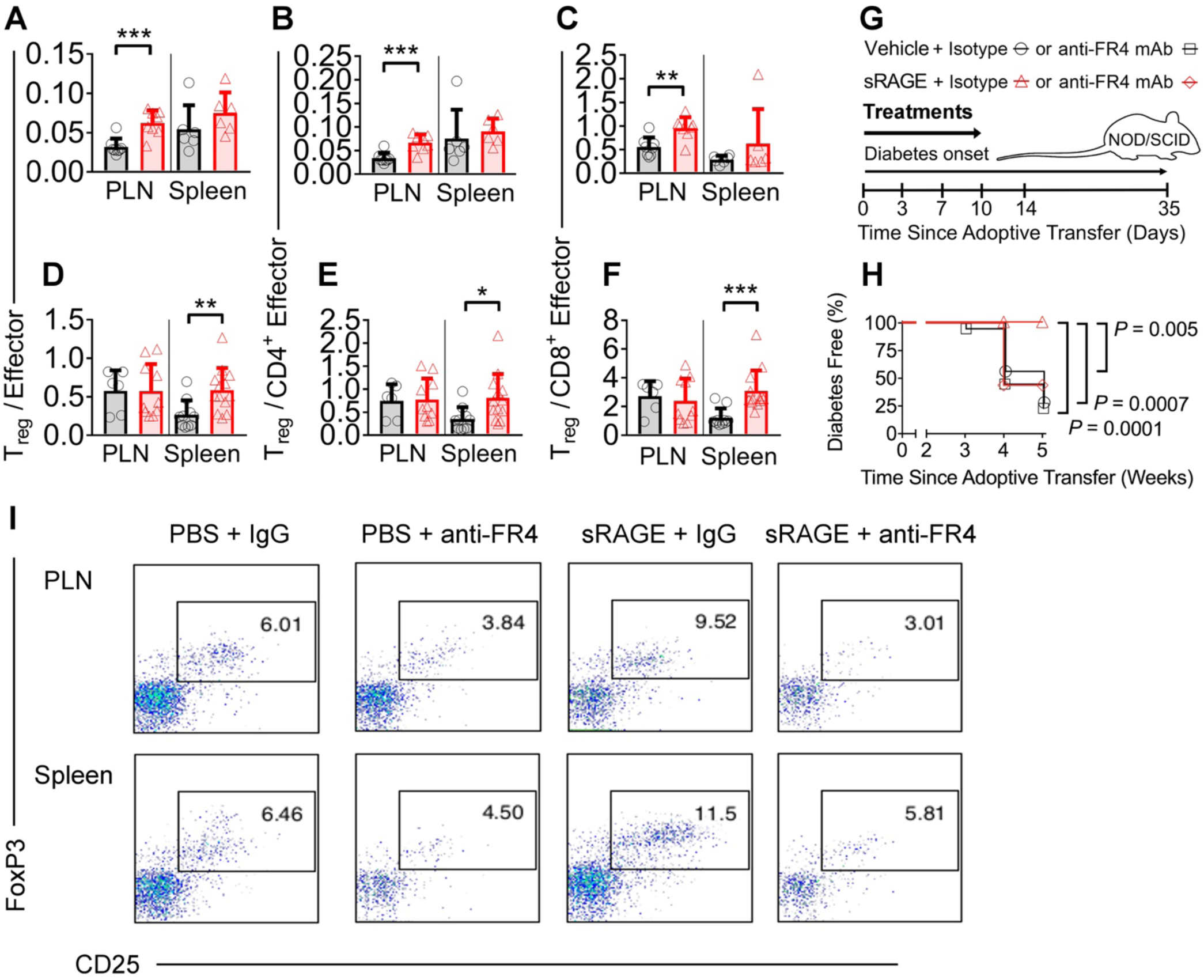
Regulatory T cells (T_regs_) are a non-redundant mechanism of action for sRAGE treatment. (A-F) Flow cytometry quantification of T_reg_/T_eff_ ratios on (A-C) day 64 and (D-F) day 225 in NOD/ShiLt mice (*n* = 4-13/group). (G) In an adoptive transfer model of autoimmune diabetes, splenocytes from diabetic NOD/ShiLt donors were adoptively transferred into NOD/SCID recipients. Recipient mice were treated with vehicle or 25 μg sRAGE twice daily for 14 days, and isotype control or anti-folate receptor 4 (FR4) antibodies at 3, 7, 10 and 14 days. (H) Diabetes incidence; and (I) Representative dot plots of CD4^+^CD8^-^CD25^+^FoxP3^+^ T_reg_ proportions (*n* = 16 mice/group). Column graphs are shown as median (IQR) and analyzed by two-tailed Mann-Whitney U-test. Diabetes incidence is shown as Kaplan-Meier survival curves and analyzed by log-rank test. \**P* < 0.05, \*\**P* < 0.01, \*\*\**P* < 0.001. PLN, pancreatic lymph nodes.

By day 225, sRAGE treated mice had decreased numbers of CD4^+^CD8^-^CD25^+^Foxp3^+^ T_regs_ and FoxP3^-^CD8^+^ T_convs_ in the PLN, as well as reduced numbers of FoxP3^-^CD4^+^ and FoxP3^-^CD8^+^ T_convs_ in the spleen, as compared with vehicle (Figure S3A-S3C). This suggested that short-term sRAGE intervention persistently dampened the immune response over the study duration. In support of this, sRAGE provided a long-lasting increase in the T_reg_/T_eff_ cell ratios in the spleen (vs. vehicle; Figure 2D-2F). Furthermore, the localized increase in T_reg_/T_eff_ cell ratios seen in the PLN of sRAGE treated mice on day 64, was resolved by day 225 (Figure 2D-2F). The proportions of CD62L^+^CD44^−^ naïve, CD62L^−^CD44^+^ effector and CD62L^+^CD44^+^ memory subsets in the T_conv_ populations remained unchanged between groups on day 225 (Figure S3B and S3C).

To determine if changes in antigen presenting cells (APCs) could influence tolerance, we also quantified the numbers of CD8^+^ and CD11b^+^ conventional dendritic cells (cDCs), plasmacytoid dendritic cells (pDCs) and macrophages on day 64. This is important because CD8^+^ cDC interactions with autoreactive CD4^+^ T cells contribute towards tolerance in NOD mice (Price et al., 2014) and, within the lymph nodes, CD8^+^ cDCs can directly induce T_regs_ via TGF-β (Yamazaki et al., 2008). CD11b^+^ cDCs and pDCs are also potent inducers of T_regs_ (Lippens et al., 2016; Tordesillas et al., 2018). Here, we found an increase in all dendritic cell subsets within the spleen but no significant changes within the PLN (Figure S4A). This supports the hypothesis that after sRAGE treatment, dendritic cells do not significantly contribute to increased proliferation or differentiation of T_regs_ within a major lymphoid tissue in T1D. Interestingly, we found increased numbers of macrophages in both the PLN and spleen (Figure S4B), consistent with previous studies using sRAGE (Pullerits et al., 2006).

Overall, these data show that sRAGE preferentially increases the numbers of T_regs_ rather than T_eff_ cells in the PLN in the short-term. In the long-term, sRAGE reduces both T_reg_ and T_eff_ cell numbers in the PLN and spleen, maintaining higher splenic T_reg_/T_eff_ ratios in mice at risk of diabetes.

### Diabetes Prevention by sRAGE is Dependent on T_regs_ and RAGE Expression

To further delineate the effects of sRAGE on T_regs_, an accelerated model of autoimmune diabetes was used that tests the immunological aspects of disease while excluding other elements such as islet dysfunction. Here, we performed the adoptive transfer of diabetes by injecting splenocytes from recently diabetic NOD/ShiLt mice into NOD/SCID recipients, that were then administered (i) vehicle and isotype control IgG, (ii) vehicle and anti-folate receptor 4 (FR4) antibodies to deplete T_regs_, (iii) sRAGE and isotype control IgG, or (iv) sRAGE and anti-FR4 antibodies to deplete T_regs_ (Figure 2G). We found that 72% and 78% of NOD/SCID recipient mice developed diabetes when given either vehicle and isotype IgG or vehicle and anti-FR4 antibodies, respectively (Figure 2H). None of the NOD/SCID mice treated with sRAGE and isotype control IgG developed diabetes, whereas 56% of those that received sRAGE and anti-FR4 antibodies to deplete T_regs_ developed diabetes (Figure 2H). We also found marked reductions in CD4^+^CD8^-^CD25^+^FoxP3^+^ T_reg_ proportions for mice treated with anti-FR4 antibodies whereas sRAGE and isotype control IgG treated mice had an increased proportion of T_regs_ (Figure 2I). These data provide direct support that T_regs_ are required for sRAGE to protect against diabetes.

sRAGE competes for RAGE ligands that are also recognized by other receptors including TLRs (Das et al., 2016). Thus, we investigated if the capacity for sRAGE to preferentially increase T_regs_ was dependent on RAGE expression. First, we established that cell surface RAGE was present on CD4^+^CD8^-^CD25^+^Foxp3^+^ T_regs_ in the PLN and spleen in NOD/ShiLt mice on day 50 of life (Figure 3A and 3B), which is when we commenced sRAGE treatment in our multi-site preclinical trial. Then, in the widely used C57BL/6 strain, we administered sRAGE to wild-type or RAGE knockout (RAGE KO) mice on days 50 to 64 of life (Figure 3C).

**Figure 3.**
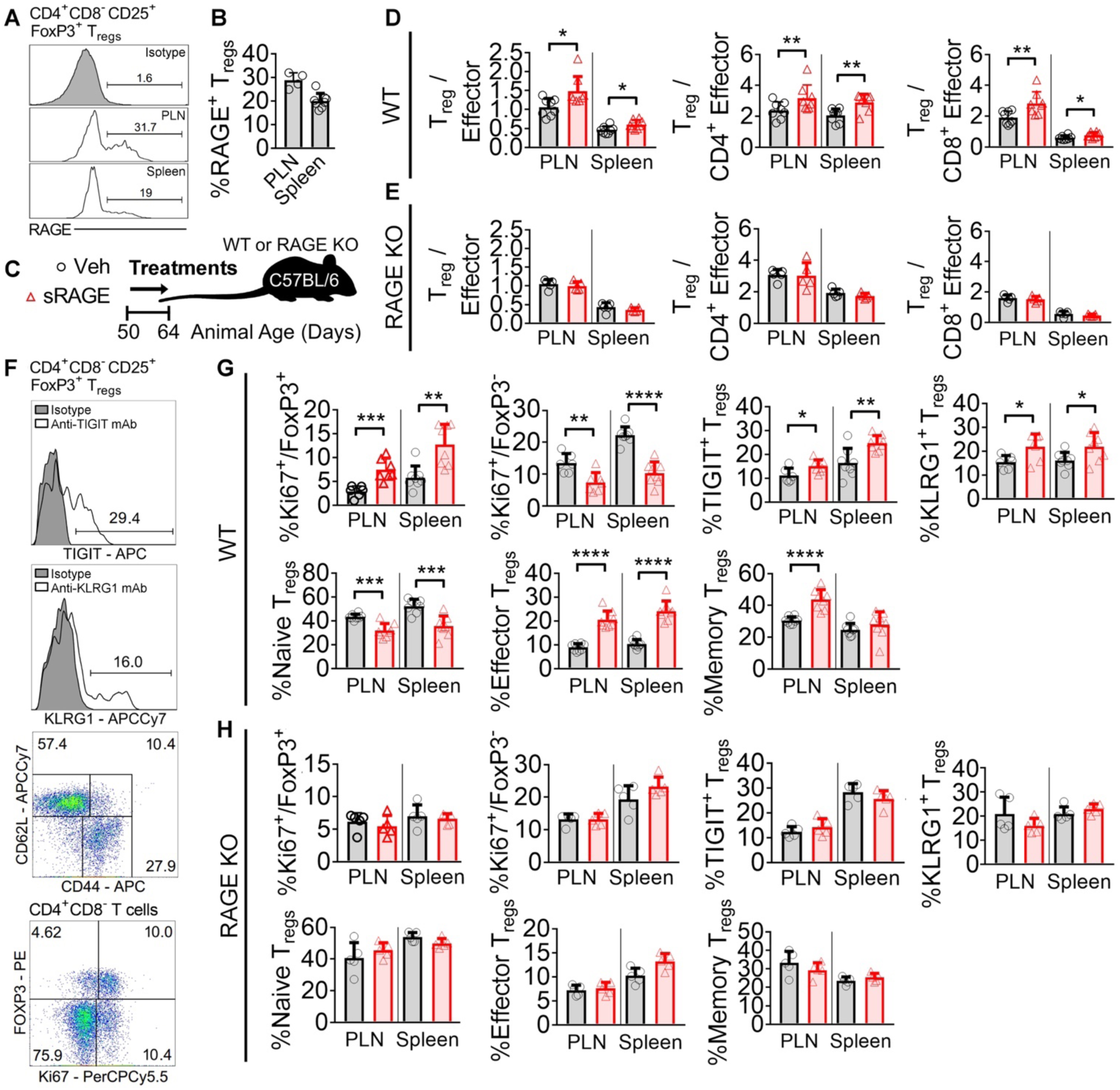
RAGE is required for the modulation of T_reg_/T_eff_ ratios by sRAGE treatment. (A and B) Flow cytometry quantification of RAGE expression on CD4^+^CD8^-^CD25^+^FoxP3^+^ T_regs_ in a mouse model of autoimmune diabetes (NOD/ShiLt) on day 50 of life. (A) Representative histograms. (B) Proportion of RAGE^+^ T_regs_ (*n* = 4-8 mice/tissue). (C) Wild-type and RAGE KO C57BL/6 mice were treated with vehicle (black bars/circles) or 25 μg sRAGE twice daily (red bars/triangles) on day 50-64 of life. (D and E) T_reg_/T_eff_ ratios on day 64 in (D) wild-type and (E) RAGE KO mice (*n* = 8 mice/group). (F-H) TIGIT, KLRG1, CD62L, CD44 and Ki67 expression. (F) Representative histograms and gating strategies. (G) Wild-type mice. (H) RAGE KO mice. Column graphs are shown as mean ± SD and analyzed by two-tailed Student’s t-test. \**P*<0.05, \*\**P* < 0.01, \*\*\**P* < 0.001, \*\*\*\**P* < 0.0001.

Consistent with our findings in NOD/ShiLt mice, there was a significant increase in the T_reg_/T_eff_ cell ratios in both the PLN and spleen of wild-type C57BL/6 mice administered sRAGE (Figure 3D). However, sRAGE treatment in RAGE KO mice did not alter T_reg_/T_eff_ cell ratios in these lymphoid tissues (Figure 3E).

We interrogated expression of the proliferation marker Ki67, as well as T_reg_ activation markers TIGIT, KLRG1, CD44 and CD62L (representative histograms and dot plots shown in Figure 3F) in the PLN and spleen of both wild-type and RAGE KO mice administered sRAGE. Consistent with the elevated T_reg_/T_eff_ cell ratios, the proportion of CD4^+^CD8^-^CD25^+^Foxp3^+^ T_regs_ expressing Ki67 in wild-type mice had increased after sRAGE treatment, whereas CD4^+^CD8^-^ Foxp3^-^ T_conv_ expression of Ki67 had declined (Figure 3G). This suggested that T_reg_ proliferation was a significant contributor to elevating the T_reg_/T_eff_ cell ratios, rather than other possible scenarios such as improved T_reg_ emigration into the lymphoid tissues. T_reg_ TIGIT and KLRG1 expression were also elevated following sRAGE administration in wild-type mice (Figure 3G), indicating the presence of activated and highly proliferative T_regs_ (Fuhrman et al., 2015). sRAGE treated T_regs_ in wild-type mice also presented a more activated phenotype based on their expression of the CD62L and CD44 adhesion molecules, with a reduced proportion of CD62L^+^CD44^−^ naïve T_regs_, and an increased proportion of CD62L^−^CD44^+^ effector and CD62L^+^CD44^+^ memory T_regs_ (Figure 3G). The expression of Ki67, TIGIT, KLRG1, CD62L and CD44 were unchanged in the RAGE KO cohort following sRAGE therapy (Figure 3H). Collectively, these data show that the activation and expansion of T_regs_ by sRAGE is dependent on cellular RAGE expression, which improves the balance of CD4^+^CD8^-^CD25^+^Foxp3^+^ T_regs_ to T_eff_ cells in mice.

### Targeted Reduction of Advanced Glycation End Products (AGEs) by sRAGE

sRAGE binds a variety of ligands implicated in the pathogenesis of T1D (Leung et al., 2016) including the heterogeneous class of compounds named advanced glycation end products (AGEs), high mobility group box 1 protein (HMGB1) and S100 calgranulins. To test if sRAGE resulted in a targeted reduction of a particular ligand, we analyzed circulating concentrations of these ligands in NOD/ShiLt mice. On day 64, immediately after sRAGE treatment, there were no changes in the circulating concentrations of AGEs, including Nε-(carboxymethyl)lysine (CML), Nε-(carboxyethyl)lysine (CEL) or methylglyoxal-derived hydroimidazolone (MG-H1; Table 1). There were also no differences in the circulating AGE precursors methylglyoxal (MGO), glyoxal (GO) or 3-deoxyglucosone (3-DG), immediately following sRAGE treatment (Table 1). However, on day 225, all plasma AGE concentrations were decreased in the sRAGE treated group (vs. vehicle; Table 1). Plasma concentrations of other RAGE ligands, S100A8, S100A9, S100B and HMGB1 did not differ between sRAGE and vehicle treated groups on either day 64 or 225 (Table 1). These findings support the hypothesis that sRAGE treatment results in a targeted long-term reduction in the circulating ligand AGEs, which is associated with protection against diabetes.

**Table 1.** Plasma concentrations of RAGE ligands in NOD/ShiLt mice. Data shown as median (IQR); two-tailed Mann-Whitney U-test; *n* = 12-15/group. \**P* < 0.05 vs. vehicle at same time point. Lys, lysine.

| Age | Day 64 |  | Day 225 |  |
| --- | --- | --- | --- | --- |
| Treatment | Vehicle | sRAGE | Vehicle | sRAGE |
| CML (nmol/mmol Lys) | 32.2 (3.5) | 31.1 (3.7) | 76.5 (10.6) | 71.1 (12.4)* |
| CEL (nmol/mmol Lys) | 13.5 (4.0) | 16.2 (8.0) | 15.4 (3.8) | 12.3 (4.2)* |
| MG-H1 (nmol/mmol Lys) | 229.3 (45.1) | 260.1 (56.4) | 267.7 (39.5) | 247.0 (54.6)* |
| MGO (nmol/L) | 499.5 (78.8) | 436.4 (142.9) | 643.7 (280.2) | 556.1 (246.4) |
| GO (nmol/L) | 848.4 (301.8) | 958.3 (316.0) | 1083.0 (505.2) | 1004.0 (518.5) |
| 3-DG (nmol/L) | 987.3 (254.7) | 1163.0 (401.0) | 2547.0 (476.0) | 2227.0 (483.0) |
| S100A8 (ng/mL) | 0.5 (0.1) | 7.1 (23.0) | 5.3 (13.1) | 5.0 (26.0) |
| S100A9 (ng/mL) | 166.8 (131.8) | 301.3 (303.0) | 74.9 (144.3) | 100.5 (81.8) |
| S100B (ng/mL) | 1081.0 (812.9) | 882.5 (512.7) | 1052.0 (528.8) | 1077.0 (545.6) |
| HMGB1 (ng/mL) | 11.9 (7.5) | 15.8 (8.2) | 7.1 (3.7) | 5.7 (5.0) |

### Islet Insulin and T_reg_ Proportions are Increased after sRAGE Treatment

We hypothesized that sRAGE could also increase the proportion of T_regs_ within pancreatic islets, thereby providing local immunoregulation and a direct improvement in insulin expression. We tested this by multiplexed immunofluorescence staining of pancreatic tissues in NOD/ShiLt mice, which showed that CD3^+^CD4^+^FoxP3^+^ T_reg_ proportions were increased on both day 64 and 225 in the sRAGE treated mice (Figure 4A and 4C). Similarly, we observed an improvement in insulin expression on both days 64 and 225 following sRAGE treatment (Figure 4B and 4C). These improvements in insulin expression were significantly correlated with the proportion of islet T_regs_ (Figure 4D), highlighting the importance of T_reg_ modulation in improving insulin expression.

**Figure 4.**
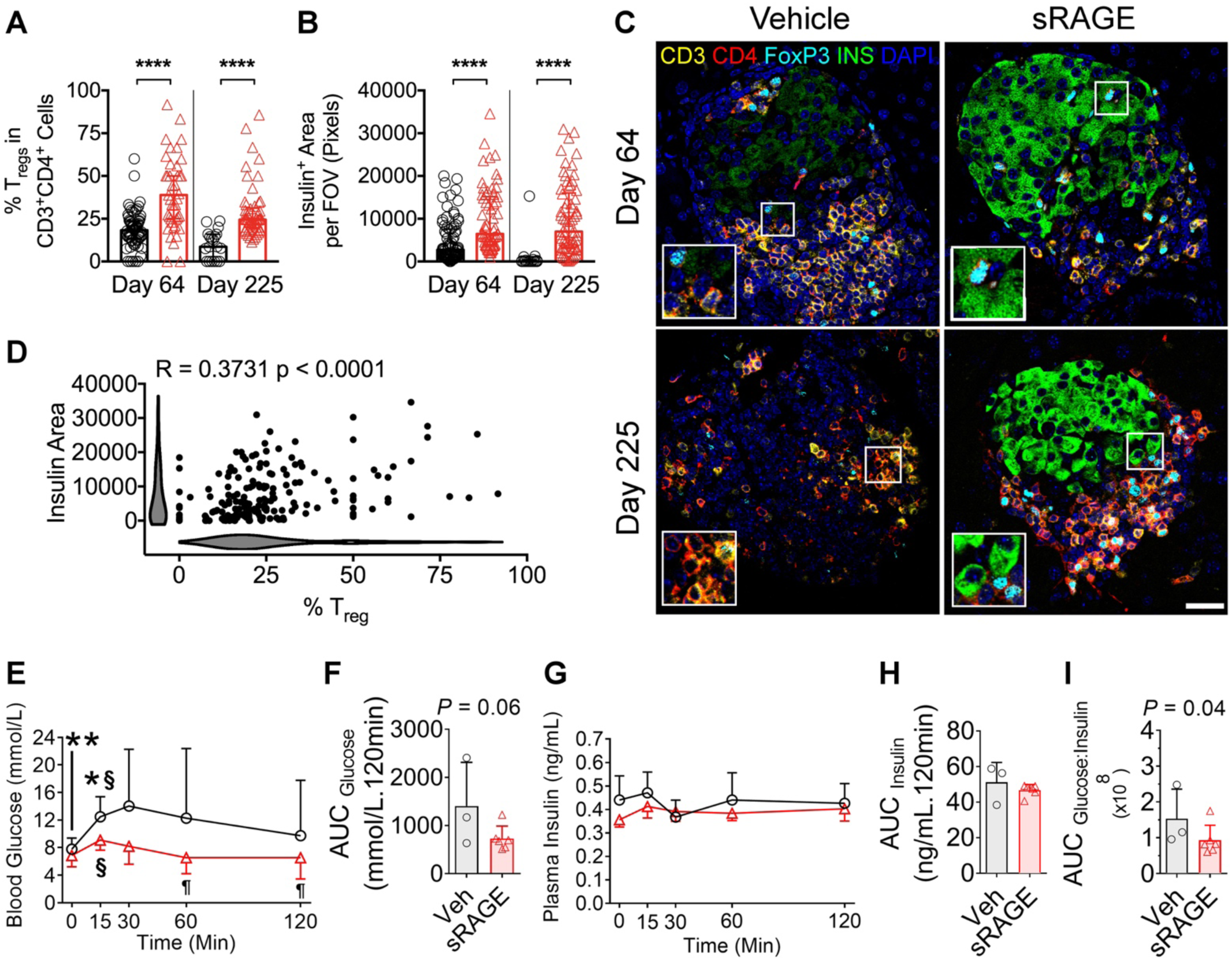
Improvements in islet infiltrating T_regs_, insulin expression and oral glucose tolerance following sRAGE treatment in NOD/ShiLt mice. (A-D) Multiplexed immunofluorescence staining and quantification of CD3, CD4, FoxP3, insulin and DAPI (*n* = 10-20 sections/mouse from *n* = 6 mice/group). Inset images are 2x magnified. (A) Proportion of islet infiltrating CD3^+^CD4^+^FoxP3^+^ T_regs_. (B) Insulin area per field of view (FOV). (C) Representative photomicrographs. Bar, 40µm. (D) Correlation analyses of CD3^+^CD4^+^FoxP3^+^ T_reg_ proportions and insulin area (shaded bars are Kernel plots showing variable distribution). (E-I) Oral glucose tolerance tests (OGTTs) on day 225 (*n* = 3-6/group). (E) Blood glucose concentrations; (F) Area under the curve for blood glucose (AUC_glucose_); (G) Plasma insulin concentrations; (H) AUC_insulin_; (I) AUC_glucose:insulin_ ratio. Data shown as mean ± SD and analyzed by two-tailed unpaired or paired Student t-tests. Correlation analyses were performed by Spearman’s test. \**P* < 0.05 between groups, \*\**P* < 0.01 between groups, \*\*\*\**P* < 0.0001 between groups. §P < 0.05 vs. 0 min within the same group, ¶P < 0.05 vs. 15 or 30 min time points within the same group.

Finally, we performed oral glucose tolerance tests (OGTTs) to assess if these improvements in insulin expression would translate into functional improvements in glucose control. The improvements in oral glucose tolerance emerged gradually, with no differences evident on day 64 (Figure S5A-S5E), an increase in insulin secretion seen prediabetes on day 80 (Figure S5F-S5J), and a marked improvement in glucose control and insulin action seen on day 225 (Figure 4E-4I). Specifically, on day 225, sRAGE treated mice had significantly lower fasting blood glucose levels before the oral glucose bolos (vs. vehicle; Figure 4E), as well as lowered blood glucose concentrations at 15, 60 and 120 minutes post-glucose bolos (vs. vehicle; Figure 4E). While the area under the curve for glucose (AUC_glucose_), plasma insulin and the AUC_insulin_ did not differ on day 225 (Figure 4F-4H), the AUC_glucose:insulin_ ratio, an indicator of insulin effectiveness, was significantly lower in sRAGE treated mice (Figure 4I), suggesting an improvement in insulin action. Altogether, these findings demonstrate that sRAGE increases T_regs_ and insulin expression in the pancreatic islets, resulting in a persisting functional improvement in oral glucose tolerance by the study end.

### sRAGE Promotes the Proliferation of Functional Human Natural T_regs_ (nT_regs_) in Co-Culture

The circulating RAGE ligand, AGEs, are independent predictors for the development of T1D in humans (Beyan et al., 2012) and sRAGE treatment led to a targeted decrease in circulating AGEs (Table 1). Therefore, we hypothesized that AGEs directly bind human T_regs_, influencing their proliferation and function. To this end, freshly isolated CD3^+^CD3^+^CD25^+^CD127^lo/-^ human T_regs_ (Figure S6A and S6B), of which 90.2 ± 7.1% were positive for FoxP3 (Figure S6C and S6D), were incubated with fluorescently labelled AGEs. These cells showed a time-dependent increase in cellular fluorescence intensity (Figure 5A), indicative of binding and cellular uptake. By contrast, co-administration of neutralizing anti-RAGE antibodies abrogated cellular fluorescence from labelled AGEs (Figure 5A), suggesting that a significant amount of AGEs bind to human T_regs_ in a RAGE-dependent manner. Human T_regs_ incubated with fluorescently labelled human serum albumin (HSA) also had less cellular fluorescence, as compared with T_regs_ incubated with AGEs, supporting the idea that AGE binding to human T_regs_ occurs is a RAGE specific mechanism (Figure 5A).

**Figure 5.**
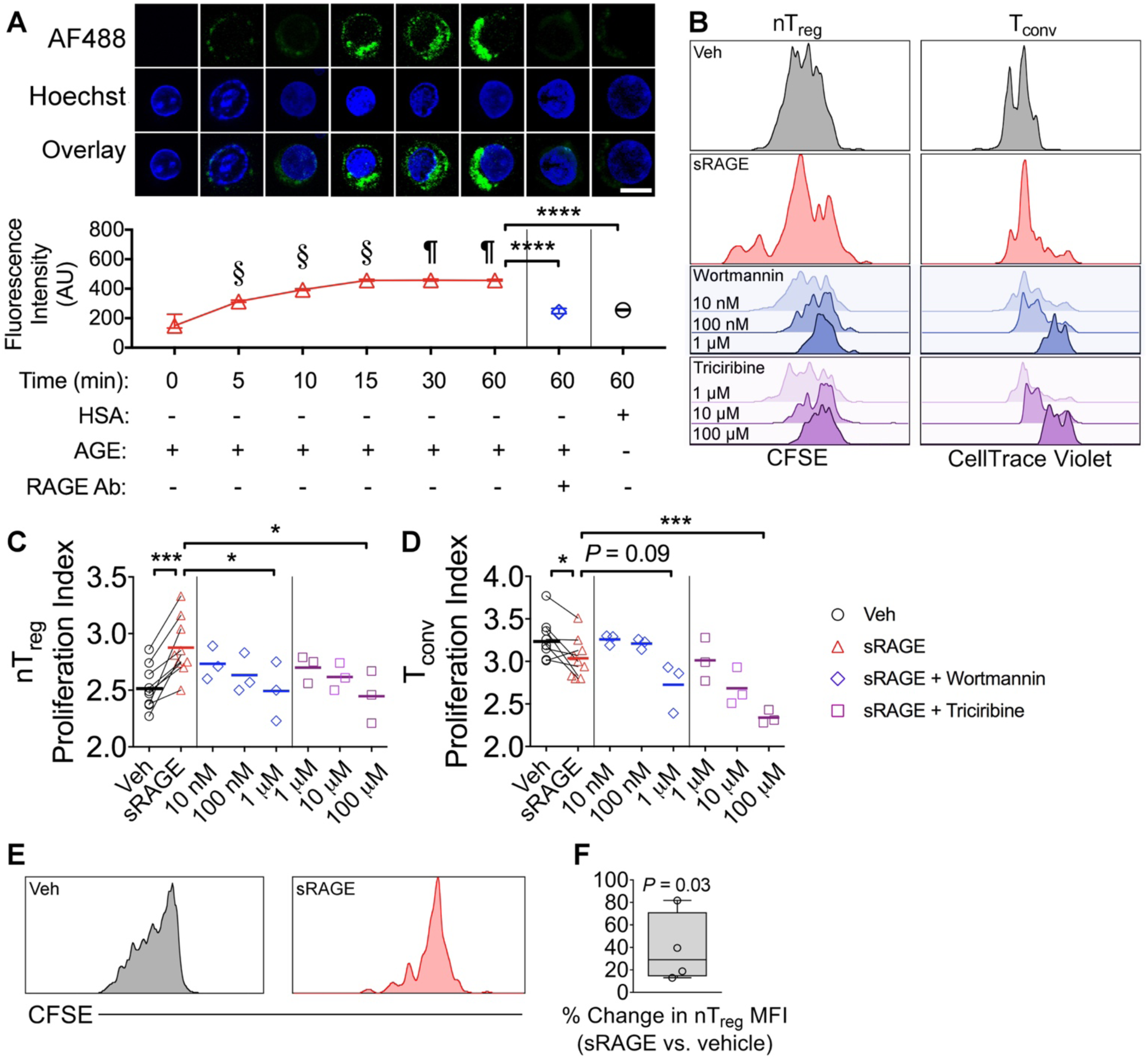
Human natural T_regs_ (nT_regs_) bind AGEs in a RAGE-dependent manner and proliferate more in co-culture when treated with sRAGE. (A) Human CD3^+^CD4^+^CD25^+^CD127^lo/–^ nT_regs_ were incubated with AGE-modified HSA (AGE-HSA or AGE) or unmodified HSA (HSA), both labelled with AlexaFluor488 (AF488). Anti-RAGE antibody was added to the culture as indicated. Bar, 10µm. *n* = 3 donors/group. (B-D) CFSE labeled CD3^+^CD4^+^CD25^+^CD127^lo/–^ nT_regs_ and CellTrace Violet labeled CD3^+^CD4^+^CD25^−^ T_convs_ were stimulated in 3 day co-culture at a 1:1 ratio containing anti-CD3/CD28 MACSiBeads (1:10, bead:cell ratio). (B) Representative histograms. (C-D) Proliferation indices of nT_regs_ and T_convs_ when administered vehicle or 50 μg sRAGE daily (*n* = 9/group), as well as sRAGE in addition to the PI3K-Akt-mTOR pathway inhibitors wortmannin or triciribine (*n* = 3/group). (E and F) CFSE labeled CD3^+^CD4^+^CD25^+^CD127^lo/–^ nT_regs_ were stimulated in 3 day monoculture containing anti-CD3/CD28 MACSiBeads (1:20, bead:cell) and 200 IU/mL IL-2. (E) Representative histograms. (F) Percent change in CFSE mean fluorescence intensity (MFI; *n* = 4/group). Data shown as mean ± SD; paired two-tailed Student’s t-tests. § P < 0.05 vs. all previous time points. ¶ P < 0.05 vs. 0, 5, 10 and 15 min. \**P* < 0.05, \*\*\**P* < 0.001. \*\*\*\**P* < 0.0001.

We tested the effects of sRAGE on various conditions conducive to the proliferation or generation of human T_regs_ *ex vivo*. First, we co-cultured CD3^+^CD4^+^CD25^+^CD127^lo/–^ natural T_regs_ (nT_regs_) and CD3^+^CD4^+^CD25^−^ T_convs_ obtained by FACS from fresh human PBMCs (Figure S6), and labeled with CFSE or CellTrace Violet respectively, in the presence of the RAGE ligand AGEs, with or without sRAGE treatment (Figure 5B). Here, we found that sRAGE accelerated nT_reg_ proliferation whilst decreasing the proliferation index of T_convs_ (Figure 5B-5D). These effects occurred through PI3K-Akt-mTOR signaling since the addition of wortmannin and triciribine, inhibitors of this pathway, resulted in dose-dependent quenching of nT_reg_ proliferation (Figure 5B-5D). This is consistent with previous findings that show the proliferation of nT_regs_ requires PI3K-Akt-mTOR activation (Wang et al., 2011). Similarly, the proliferation of sRAGE treated T_conv_ cells was further decreased in the presence of wortmannin and triciribine (Figure 5B-5D). Altogether, this data supports a role for sRAGE in promoting nT_reg_ proliferation in co-culture.

We also tested if sRAGE could promote the generation of induced T_regs_ (iT_regs_). Here, CD3^+^CD4^+^CD45RA^+^CD45RO^−^ naїve human T cells were isolated (Figure S7A and S7B), stimulated as previously described (Ellis et al., 2012) in the presence of AGEs and analyzed for iT_reg_ differentiation (Figure S7C-S7E). In these conditions, sRAGE treatment led to a reduction in iT_reg_ generation (Figure S8), which suggests the differentiation of iT_regs_ does not contribute to increasing the overall population of T_regs_ after sRAGE treatment.

### AGEs Promote Human nT_reg_ Proliferation in Monoculture but Inhibit Suppressive Function

To further investigate the hypothesis that sRAGE maintains the proliferative balance between nT_regs_ and T_convs_, we performed nT_reg_ monoculture experiments in the presence of AGEs with or without sRAGE administration (Figure 5E). Here, we found that sRAGE increased the mean fluorescence intensity (MFI) of CFSE labeled nT_regs_ (vs. vehicle; Figure 5E and 5F), suggesting that sRAGE inhibits the proliferation of nT_regs_ when other T cell populations are absent. This finding is consistent with previous studies where RAGE KO inhibits T_conv_ proliferation (Chen et al., 2008). To the best of our knowledge, however, there are no studies showing a role for RAGE in nT_reg_ proliferation. Here, our data supports a role for the RAGE ligand, AGEs, in promoting the proliferation of nT_regs_ cultured in isolation.

However, proliferation of nT_regs_ can be associated with a loss of suppressive function (Gerriets et al., 2016; Putnam et al., 2009), so we postulated that although AGEs induce the proliferation of nT_regs_ in the absence of other T cell populations, they limit their function. To test this, we performed human nT_reg_ monoculture experiments and quantified changes in gene expression using NanoString. Principal component analyses (PCA) distinctively separated the HSA (control) and AGE treated nT_regs_ (Figure 6A), particularly along principal component 1 (PC1) that accounted for 56% of the variance overall. HSA treated nT_regs_ were enriched in the expression of genes involved in nT_reg_ function (*JAK1*, *IRF4*, *SOCS1*; Figure 6A) (Cretney et al., 2011; Kirken et al., 1995; Takahashi et al., 2017), while AGE treatment decreased the expression of these functional nT_reg_ genes.

**Figure 6.**
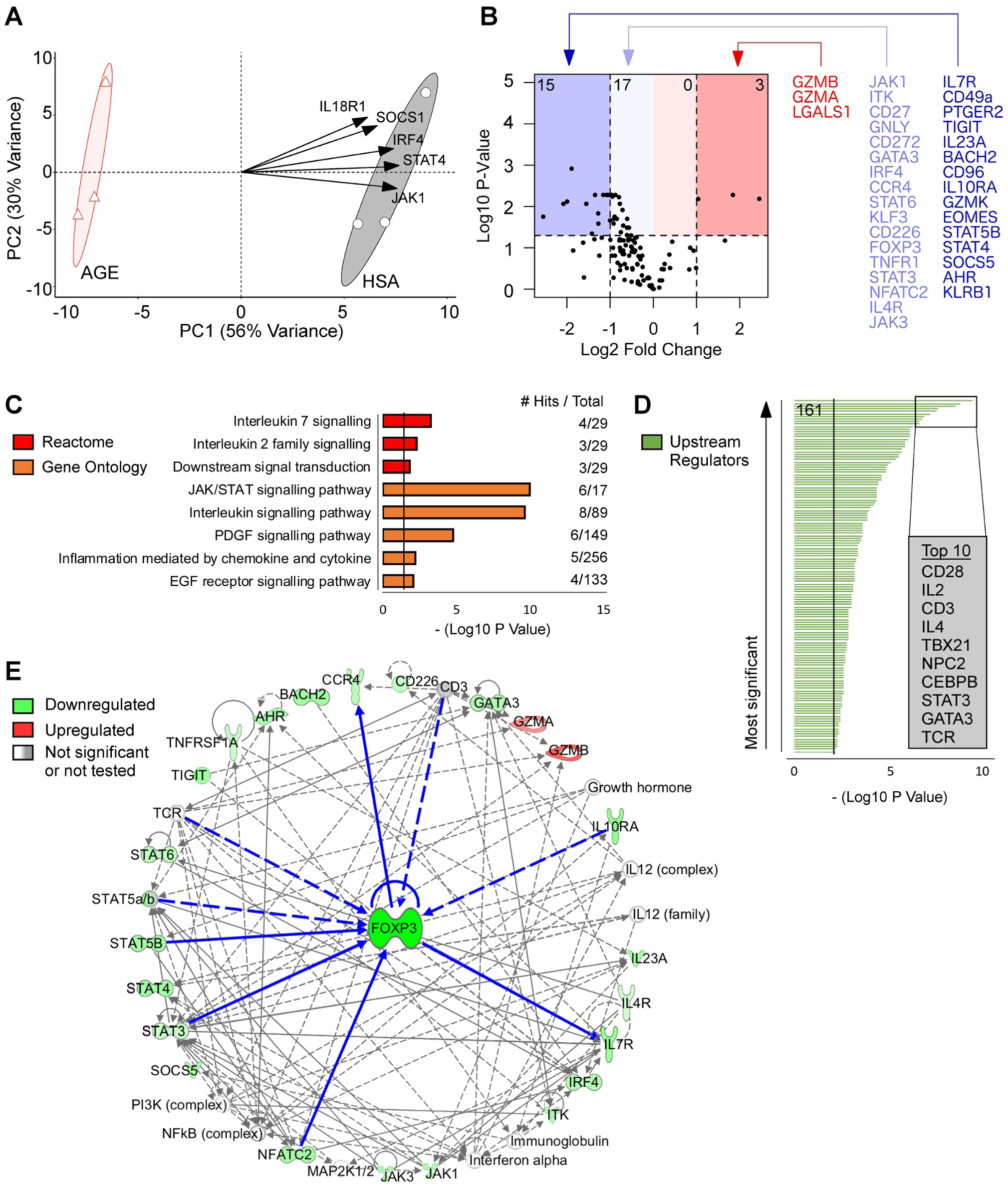
AGE treatment of human nT_regs_ downregulates key genes for suppressive function and manipulates signaling cascades upstream and downstream of *FOXP3*. (A) Principal component analysis (PCA) with the top five variable loadings shown as arrows. (B) Volcano plot visualization for changes in gene expression (number of differentially expressed genes shown in top corners). (C) Pathway enrichment analysis using the Reactome and Gene Ontology databases. (D) Upstream regulator predictions using Ingenuity Pathway Analysis (161 significant regulators were identified). (E) Network analysis using *FOXP3* as the focus node. Neighboring molecules are connected to *FOXP3* with bolded blue lines. All other molecules are 2-3 nodes adjacent to *FOXP3*. Direct relationships are shown as solid lines, indirect relationships are shown as dashed lines. *n* = 3/group, *P* values were corrected by false discovery rate or Bonferroni adjustment.

Differentially expressed genes were visualized by volcano plot (Figure 6B) where we found that AGE treated nT_regs_ had reduced expression of key nT_reg_ genes including *FOXP3*, *IL7R*, *TIGIT* and *STAT5b* (Cohen et al., 2006), as well as several other genes that promote nT_reg_ function (*BACH2*, *CD96*, *IL10RA, JAK1/3, ITK, CD27, IRF4, STAT3/6, CD226*) (Chaudhry et al., 2011; Cretney et al., 2011; Duggleby et al., 2007; Fuhrman et al., 2015; Huang et al., 2014; Kirken et al., 1995; Pallandre et al., 2007; Piedavent-Salomon et al., 2015; Roychoudhuri et al., 2013; Sanchez-Guajardo et al., 2007) and migration (*CCR4*, *AHR*) (Gobert et al., 2009; Ye et al., 2017). Interestingly, AGE treated nT_regs_ also had changes in the expression of granzymes A/B/K and granulysin (*GZMB*, *GZMB*, *GZMK, GNLY*) and decreased expression of *EOMES* (Figure 6B), a marker for T cell exhaustion.

Pathway enrichment analysis of genes downregulated by AGEs identified significant biological pathways including interleukin 7 and interleukin 2 family signaling, JAK/STAT signaling pathway, as well as the PDGF/EGFR signaling pathways (Figure 6C) that all play a role in maintaining nT_reg_ function (Lo Re et al., 2011; Wang et al., 2016). Ingenuity Pathway Analysis predicted 161 statistically significant upstream regulators associated with the changes in gene expression caused by AGE treatment (Figure 6D and Table S1). Of interest, the top ten upstream regulators included key nT_reg_ signaling molecules (*CD28*, *IL2*, *CD3*, *STAT3*, TCR) as well as other T cell associated proteins (*IL4*, *TBX21*, *NPC2*, *CEBPB*, *GATA3*; Figure 6D). These data are consistent with a role for the RAGE ligand, AGEs, in influencing crucial nT_reg_ signaling cascades. Finally, we performed network analysis, focusing on the most statistically significant network (Table S2), which identified neighboring nodes for *FOXP3* and several genes upstream (*IL10RA*, *STAT3/5B*, *NFATC2*) and downstream (*IL7R*, *CCR4*) of *FOXP3*, which were changed in expression following AGE treatment (Figure 6E). Predicted upstream regulators after AGE treatment (*CD3*, TCR; Figure 6D) were also neighboring nodes with *FOXP3* (Figure 6E). All other genes were either two or three nodes adjacent of *FOXP3* (Figure 6E), highlighting the intimate cross-talk amongst genes influenced by AGEs and the nT_reg_ master regulatory gene *FOXP3*.

These data provide strong evidence that despite promoting human nT_reg_ proliferation in the absence of other T cell populations, AGE treatment results in a loss of suppressive function. These functional defects can be rescued by the competitive RAGE antagonist sRAGE in co-culture where it expands functional human nT_regs_ that suppress T_conv_ proliferation. Taken together with the efficacy of sRAGE to expand T_regs_ and prevent diabetes in mice, we propose that sRAGE therapy should be tested in at-risk individuals to prevent the onset of T1D.

## Discussion

There is an urgent unmet need for novel disease targets and treatments that are reproducible and have high potential to be translated to prevent T1D in humans (Insel et al., 2015). Here, we report that a RAGE antagonist sRAGE has a novel immunomodulatory role where it maintains the balance between T_regs_ and T_effs_, a critical process in self-tolerance (Brusko et al., 2005). We show that sRAGE treatment in several mouse models resulted in increased T_reg_ ratios in the PLN and spleen, which are lymphoid tissues that have both been implicated in the pathogenesis of diabetes (McNally et al., 2011), as well as within the pancreatic islet immune cell infiltrate. Using human T cell culture, we found that sRAGE promoted the expansion of functional human nT_regs_, whereas the RAGE ligand AGEs significantly impaired their suppressive function. Ultimately, we show that short-term intervention with sRAGE restored T_reg_/T_eff_ ratios, protecting against diabetes onset at two independent research centers.

Immediately after sRAGE therapy on day 64, islet infiltration was reduced, which was accompanied by increases in the absolute numbers of T_regs_ as well as T_reg_/T_eff_ ratios in the PLN. These changes in the PLN are critical for islet preservation and arresting disease progression, since T_regs_ localized here suppress the priming of autoantigen specific T cells (McNally et al., 2011), which would otherwise infiltrate the pancreatic islets. The transient increase in PLN leukocytes is consistent with previous work where sRAGE showed immunomodulatory effects on monocytes, macrophages and B cells (Pullerits et al., 2006). Here, we also found an increase in macrophage numbers following sRAGE treatment on day 64. Further investigation into their phenotype is warranted since changes in the macrophage transcriptome associates with T1D (Ferris et al., 2017). Interestingly, we report increased numbers of dendritic cell subsets within the spleen, but not the PLN, suggesting that they are not significant contributors in the expansion of T_regs_ within the major draining lymphoid tissue in T1D. In support of this, we also delineated that the modulation of T_regs_ specifically, using anti-FR4 antibodies to deplete T_regs_, is critical for the protection afforded against diabetes by sRAGE. T_reg_ immunomodulation comparable to this has also been seen with other interventions that are lead candidates for the treatment of T1D (Sherry et al., 2011). Further development of sRAGE therapy to optimize the dosing regimen and therapeutic window is likely to result in even more efficacious outcomes, which will facilitate its competition with the current lead candidate agents. We also acknowledge that recombinant human sRAGE is unlikely to be immunogenic since it is an endogenous protein, but it would be worthwhile investigating if RAGE-specific autoantibodies were produced as a result of this treatment.

By day 80, islet infiltration was modestly increased in sRAGE treated mice, which could reflect differences in the type of infiltrating immune cells present. Indeed, it is likely that there are increased numbers of infiltrating T_regs_ at this time point in sRAGE treated mice, but we highlight that this remains untested. In support of this hypothesis, we found that sRAGE treatment had increased the proportion of T_regs_ in the islets on days 64 and 225, which associated with reduced islet infiltration by the study end. By day 225, sRAGE treatment had also improved oral glucose tolerance and insulin expression to preserve pancreatic islet numbers, which reflects the protection against diabetes (Sosenko et al., 2012). Given that early short-term intervention with sRAGE improved the balance of T_regs_, imparting persistent improvements in autoimmunity and glycemic control, this therapy shows promise in preventing the onset of clinical T1D.

Mice deficient in RAGE could not increase T_reg_/T_eff_ cell ratios within lymphoid tissues following sRAGE intervention, which alludes to the importance of cellular RAGE antagonism for the mechanism of this therapy. RAGE plays an important role in T cell survival (Durning et al., 2016) and proliferation (Chen et al., 2008), including in individuals at-risk of T1D, but there has been no delineation as to whether this applies to specific T cell subsets such as T_regs_. Furthermore, there is no data examining whether RAGE impacts T_reg_ function. In the current study, we confirmed the expression of RAGE on T_regs_ in the PLN and spleen of NOD/ShiLt mice and found that sRAGE treatment improved localized T_reg_/T_eff_ ratios after therapy completion. In the long-term, sRAGE treated mice had reduced circulating concentrations of the heterogeneous class of RAGE ligands termed AGEs, including CML, CEL and MG-H1, and increased splenic T_reg_/T_eff_ cell ratios. Similarly, sRAGE increased T_reg_/T_eff_ ratios, as well as the expression of the T_reg_ activation markers Ki67, TIGIT, KLRG1 and CD44, in wild-type C57BL/6 but not RAGE KO mice. It was noteworthy that RAGE KO mice did not have changes in T_reg_/T_eff_ ratios as compared with WT mice prior to sRAGE treatment. This could suggest that long-term depletion of RAGE leads to the normalization of T_reg_/T_eff_ ratios, potentially by compensatory signaling through receptors which share ligands with RAGE such as toll-like receptors (TLRs). In support of this, crosstalk between RAGE and TLRs has been found to synergistically drive inflammation via NF-κB (Gasiorowski et al., 2018). However, as alluded to in this study, the function of T cells and T_regs_ in RAGE KO mice may be compromised.

In children who progress to T1D, circulating sRAGE concentrations are decreased prediabetes (Forbes et al., 2011; Salonen et al., 2015; Salonen et al., 2016) and increases in the RAGE ligand AGEs are independent predictors of progression to T1D (Beyan et al., 2012). To this end, short-term sRAGE treatment facilitated long-lived decreases in circulating protein-bound AGEs (CML, CEL and MG-H1), alongside improved insulin expression and action in mice. We also demonstrated that AGEs promoted human nT_reg_ proliferation when these cells were cultured alone but this resulted in significant impairment of nT_reg_ function. By contrast, sRAGE treatment led to the expansion of functional human nT_regs_, since sRAGE increased the proliferation of nT_regs_ that could inhibit T_conv_ cell division in co-culture.

We highlight the importance for future studies to perform the functional assays presented in this study not only using healthy human T_regs_, but also in T cells obtained from individuals with presymptomatic and overt T1D, to expand our knowledge about the role of RAGE in T_reg_ function in this disease context. Furthermore, the use of tetramer complexes could help confirm if the effects observed on T cells in this study translate to changes in the numbers and function of autoantigen-specific T cells and T_regs_ that are known to affect T1D pathogenesis. Finally, the use of humanized mice with human umbilical cord blood stem cells or bone marrow, liver and thymus (“BLT”) transplants could further progress the therapeutic development of sRAGE by demonstrating its efficacy using an *in vivo* human immune system. Overall, RAGE and RAGE ligand biology clearly require further interrogation to better understand their contribution to antigen-specific T cell and T_reg_ responses in T1D.

In summary, we have shown that sRAGE is an immunomodulatory compound and is an important factor in balancing T_reg_ responses to their environment, including through the RAGE ligand AGEs. This was ascertained using human T cell assays as well as using murine models that were RAGE deficient or at high-risk of autoimmune diabetes, where the presence of functional T_regs_ is known to be compromised. We showed that a short-term two-week intervention with sRAGE elicited persistent effects on the immune system, ultimately preserving β cell function and reducing diabetes incidence in a multi-site preclinical trial. Given that sRAGE is a native protein that is reduced in children at risk of T1D, and has persistent benefits on insulin expression, oral glucose tolerance and T_reg_/T_eff_ ratios, we suggest that this treatment is a promising new therapy for further development to prevent T1D.

## Materials and Methods

### Mice

Female NOD/ShiLt mice were housed in specific pathogen-free conditions at two independent research centers – Site 1, Translational Research Institute, Brisbane, Australia; and Site 2, Type 1 Diabetes Research Center, Novo Nordisk, Seattle, US. Mice were sourced from The Animal Resources Centre (Canning Vale, Australia; Site 1) or The Jackson Laboratory (Sacramento, CA, US; Site 2), and were provided free access to irradiated diet (Site 1, Specialty Feeds Rat and Mouse Diet; Site 2, Purina Lab Diet 5053) and water (Site 1, autoclaved; Site 2, filtered). Sample size calculations were performed using α = 0.05 and a power of 0.80, where 80% of the control mice were expected to develop diabetes by the study end as per historical data available in the animal facilities.

Female NOD/SCIDs, wild-type C57BL/6 and RAGE KO C57BL/6 (Liliensiek et al., 2004) mice were housed at Site 1. Animal studies were approved at both sites by their respective institutional ethics committees and adhered to national guidelines by the National Health and Medical Research Council (NHMRC, Australia) and NIH (US).

### Human Samples

Human whole blood specimens were obtained by venipuncture using citrate-phosphate-dextrose-adenine or EDTA anti-coagulants and transported to the laboratory within 1-2 hours. Human donors were healthy volunteers, 18-65 years of age and provided informed consent. Experiments were approved by the Mater Ethics Committee.

### sRAGE Therapy

Randomized mice were intraperitoneally injected on days 50-64 of life with 100 µL recombinant human sRAGE (25 µg) twice daily (Sites 1 and 2), vehicle (PBS) twice daily (Site 1), sRAGE (100 µg) once daily (Site 2), or untreated (Site 2). Treatment dosages was based on previous studies using sRAGE, which were able to detect changes in end points using 25-100 µg sRAGE per day (Pullerits et al., 2006; Wang et al., 2010). Recombinant sRAGE was produced using an insect cell and baculovirus expression system with the cloned sequence for human endogenous secretory RAGE (esRAGE; from herein, sRAGE). Recombinant sRAGE was isolated using size exclusion and affinity chromatography, and protein purity determined to be >99% by SDS-PAGE and Western Blot. Endotoxin levels were 0.065 EU/mg (0.00325 EU/day and 0.0065 EU/day for 25 µg twice daily and 100 µg once daily dosages, respectively) as determined by limulus amebocyte lysate (LAL) assay.

Mice were fasted for 4-6 h and euthanazed on day 64, 80 or for non-progressors on 225 of life. Non-fasted blood glucose concentrations were measured weekly using a glucometer (Site 1, SensoCard; Site 2, Bayer Contour USB) between days 50-225. Diabetes was diagnosed when this exceeded 15 mmol/L on consecutive days at which point these progressors left the study.

### Adoptive Transfer

Splenocytes (10^7^) from untreated female NOD/ShiLt mice diagnosed with diabetes within seven days were intravenously injected (200µL) into 5-9 week old female NOD/SCID recipients (Leiter, 2001). Briefly, splenocytes were mechanically dissociated using 70µm filters, red cell lysis performed using ammonium-chloride-potassium (ACK) buffer (ThermoFisher) and splenocytes washed several times into serum-free mouse-tonicity PBS (MT-PBS) for injection *in vivo*. Randomized NOD/SCID mice were then administered the following treatments for two weeks post-adoptive transfer: (i) PBS and rat isotype control IgG2b antibodies (RTK4530; BioLegend), (ii) PBS and anti-FR4 antibodies (TH6; BioLegend) for the depletion of T_regs_ via their selective expression of FR4 (Yamaguchi et al., 2007), (iii) sRAGE and isotype control antibodies, or (iv) sRAGE and anti-FR4 antibodies. PBS and sRAGE (25 µg) were given twice daily, as above. Isotype control and anti-FR4 antibodies (10 µg for both) were given on days 0, 3, 7, 10 and 14, and endotoxin levels were <0.01 EU/µg (<0.1 EU/injection) as determined by LAL assay. Diabetes was monitored and diagnosed as above.

### Oral Glucose Tolerance Tests (OGTTs)

Mice were fasted for 4-6 h and administered a 2 g/kg glucose bolus by intragastric gavage. At 0, 15, 30, 60 and 120 min post-glucose bolus, blood glucose and plasma insulin were measured by glucometer and ELISA (Crystal Chem), respectively.

### RAGE Ligand Assays

Fasting plasma S100A8/A9 (R&D DuoSet), S100B (Abbexa), and HMGB1 (Shino-Test) were measured by ELISA. Circulating AGEs and dicarbonyls were measured by liquid chromatography-tandem mass spectrometry (LC-MS/MS) as previously described (Scheijen and Schalkwijk, 2014).

### Islet Histology and Immunofluorescence Staining

Formalin fixed tissue sections (4-5µm) were deparaffinized and rehydrated using standard techniques, then stained with H&E, coverslipped in DPX and imaged on the VS1200 brightfield microscope (Olympus). Islet infiltration was assessed in a blinded fashion using an islet infiltration index from 0 to 1 (no infiltration to complete infiltration), and individual islets graded as 0, 1, 2, 3 or 4 (no infiltration to >75% infiltration) as previously described.

For multiplexed immunofluorescence staining, antigen retrieval was performed using sodium citrate buffer (pH 6). Non-specific binding was blocked using 10% donkey serum at room temperature for 1 hour, then sections were incubated with rabbit anti-CD3 (SP7, Abcam), goat anti-CD4 (#AF554, polyclonal; R&D Systems) and biotinylated anti-FoxP3 (FJK-16s; eBioscience) antibodies overnight at 4°C, followed by incubation with anti-rabbit IRDye 800CW (#926-32213; Li-cor Biosciences), anti-goat AlexaFluor568 (#A11057; ThermoFisher), and streptavidin-AlexaFluor647 (#S32357; ThermoFisher) at room temperature for 1 hour. Sections were blocked again as described above, then incubated with rat anti-insulin antibody (182410; R&D Systems) overnight at 4°C, followed by incubation with anti-rat AlexaFluor488 (#A21208) at room temperature for 1 hour. Sections were coverslipped in Fluoroshield with DAPI and images captured on the FV1200 confocal microscope (Olympus). Quantification was performed in a blinded fashion using ImageJ v2.0.0.

### Flow Cytometry and Cell Sorting

Mouse spleen and PLN were mechanically dissociated into single cells using 40µm filters and red cell lysis performed using ACK buffer (ThermoFisher). Blocking was performed using anti-CD16/CD32 antibodies (BD Biosciences) and cells were stained using antibodies against CD4 (RM4-5, BD Biosciences unless indicated), CD8 (53-6.7), CD11b (M1/70), CD11c (HL3), B220 (RA3-6B2), F4/80 (CI:A3-1; Bio-Rad), CD62L (MEL-14), CD44 (IM7), CD25 (PC61), FoxP3 (FJK-16s, eBioscience), TIGIT (1G9, BioLegend unless indicated), KLRG1 (2F1) and Ki67 (16A8). Samples were analyzed on the LSRII (BD Biosciences) and FlowJo (Tree Star Inc).

Human PBMCs were isolated by Ficoll and incubated with antibodies against CD3 (OKT3, BioLegend unless indicated), CD4 (RPA-T4), CD25 (BC96), CD127 (A019D5) and FoxP3 (PCH101; eBioscience). Dead cells were excluded using a Live/Dead viability dye (ThermoFisher) and blocking was performed using Human TruStain FcX (BioLegend). CD3^+^CD4^+^CD25^+^CD127^lo/-^ nT_regs_ and CD3^+^CD4^+^CD25^-^ T_convs_ were isolated on the Astrios (Beckman Coulter) or FACSAria (BD Biosciences) and analyzed using FlowJo (>97% nT_reg_ and T_conv_ purity). CD3^+^CD4^+^CD25^+^CD127^lo/-^ nT_regs_ were 90.2 ± 7.1% FoxP3 positive using human PBMCs (Figure S6C and S6D), consistent with previous studies which have examined the use of CD3^+^CD4^+^CD25^+^CD127^lo/-^ cells for isolating live, viable nT_regs_ for *ex vivo* assays (Putnam et al., 2009).

### Human T Cell Culture

Human nT_regs_ were CFSE labelled and cultured in TexMACs (Miltenyi Biotec) supplemented with 10% heat inactivated fetal bovine serum (HI-FBS, Life Technologies unless indicated), 100 U/mL penicillin-streptomycin, 10 µM β-mercaptoethanol and 100 µg/mL AGE-modified HSA (AGE-HSA or AGE) at a final number of 2.5×10^4^ cells in a U-bottom 96-well plate. T_convs_ were CellTrace Violet labelled and added to a final number of 2.5×10^4^ cells. Cells were stimulated using anti-CD3/CD28 MACSiBeads at a 1:10 bead-to-cell ratio. Co-cultures were treated with 50 µg sRAGE or PBS daily, for 72 h at 37 ^O^C/5% CO_2_. Proliferation was analyzed using CFSE and CellTrace Violet dye dilution on the LSRFortessa (BD Biosciences) and FlowJo.

CFSE labeled human nT_regs_ were also grown in monoculture using TexMACs supplemented with 10% HI-FBS, 100 U/mL penicillin-streptomycin, 10 µM β-mercaptoethanol and 200 IU/mL IL-2 at a final number of 2×10^4^ cells in a U-bottom 96-well plate. Cells were stimulated using anti-CD3/CD28 MACSiBeads at a 1:20 bead-to-cell ratio. Cells were treated with 100 ug/mL HSA or AGEs for 72 h at 37 ^O^C/5% CO_2_, and analyzed as above.

For gene expression analysis, unlabeled nT_regs_ were grown in monoculture under the conditions as described above for 72 h and anti-CD3/CD28 MACSiBeads removed by EasySep Magnet. RNA was isolated using RNAzol RT (Astral Scientific) per manufacturer’s instructions with the use of molecular grade isopropanol and ethanol (Sigma Aldrich) for RNA precipitation. Briefly, cells were lysed and centrifuged to separate the aqueous phase which contained RNA.

Multiple ethanol washes were performed before resuspension of the purified RNA pellet. RNA quality and quantity were probed using an Implen NanoPhotometer N60 (LabGear). 100 ng RNA was hybridized overnight using a NanoString Custom CodeSet for 136 T cell specific genes. Normalization of raw counts was performed using NanoString NSolver software with the housekeeping genes *ACTB*, *B2M*, *GAPDH*, *HPRT1* and *RPLP0* (panel in Table S3).

Naïve CD4^+^ T cells were negatively isolated from fresh PBMCs by EasySep (Stem Cell Technologies; >95% purity). Cells were stimulated (Ellis et al., 2012), treated with 100 µg/mL AGEs and 50 µg sRAGE or PBS daily for 72 h at 37 ^O^C/5% CO_2,_ and analyzed for CD3^+^CD4^+^CD25^+^CD127^lo/–^ iT_reg_ differentiation on the LSRFortessa (BD Biosciences) and FlowJo.

### AGE-HSA Production

AGE-HSA was produced by incubating 20 mg/mL fatty acid-free, cold ethanol precipitated human serum albumin (Sigma) and 0.5 M D-(+)-glucose (Sigma) in HyClone PBS (GE Healthcare) for 3 months at 37°C in the dark. Solutions were dialyzed in PBS (GE Healthcare) with 10K MWCO Slide-A-Lyzer Cassettes (ThermoFisher), 0.22 µm filtered and stored at −80°C.

Endotoxins were determined to be <1 EU/mL (<0.005 EU/mL at final concentrations in human cell culture experiments) by Limulus Amebocyte Lysate Assay. Quantification of AGE-HSA glycation adducts – RAGE ligands – was performed using LC-MS/MS as described above, producing the following concentrations: 646,568 nM CML, 45,279 nM CEL and 198,454 nM MG-H1 (28-, 22- and 11-fold increases in glycation adduct concentrations respectively, as compared with unmodified HSA).

### Human nT_reg_ Binding Assay

Human nT_regs_ were stained using Hoechst 33342 (ThermoFisher) and cultured in phenol-free RPMI-1640 (ThermoFisher) in glass chambers and administered 100 µg/mL HSA-AlexaFluor488, AGE-AlexaFluor488, or AGE-AlexaFluor488 and anti-RAGE antibody (#AB5484, Merck) for the durations indicated. Cells were imaged on the FV1200 confocal microscope and quantification was performed in a blinded fashion using ImageJ.

### Quantification and Statistical Analysis

Statistical analyses were performed using GraphPad v5.01 and *P* < 0.05 was considered statistically significant. Comparisons were done using biological not technical replicates which are shown in all figures. Normality was tested by Kolmogorov-Smirnov test. Means were compared by two-tailed Student’s t-test and shown as mean ± SD. Medians were compared by two-tailed Mann-Whitney U-test and shown as median (IQR). Kaplan-Meier survival curves were compared by log-rank test. Regression lines were compared by ANCOVA. Proportions were compared by Fischer’s test. Gene expression was analyzed using R v.3.4.4 for PCA and volcano plots, DAVID v.6.8 for Reactome Pathway enrichment, PANTHER v.14.0 for Gene Ontology overrepresentation and Ingenuity Pathway Analysis v.1.14 for identification of upstream regulators and network analysis (*P* values were adjusted using false discovery rate or Bonferroni).

## Author Contributions

SSL designed and performed experiments, analyzed and interpreted data and prepared the manuscript. DJB, DAMc, AZ, AKF, NF, TW, JJM, JLS and CGS performed experiments. TEB designed and JC performed the incidence experiment at the second site and both completed its data analyses and interpretation. KJR assisted with the human study design and completion. RJS, P-HG, MK, led by JMF conceptualized and designed the overall study, gained financial support and completed data analyses and interpretation. All authors edited and approved the final manuscript.

## Acknowledgments

The authors thank I. Rojas and S. Diaz-Guilas for assistance with human cell culture; C. O’Brien and T. Friesen for assistance with the incidence study at Site 2; I. Buckle, J. Naranjo and E. Hamilton-Williams for providing diabetic NODs for the adoptive transfer; J. Lynch and S. Phipps for providing RAGE KO mice for exploratory analyses; K. MacDonald for feedback on T_reg_ experiments; E. Williams and the Australian Equine Genetics Research Centre for assistance with the RAGE KO mice; and the Biological Resources, Flow Cytometry, Histology and Microscopy facilities at the Translational Research Institute.

## Supplemental Figures Titles and Legends

**Figure S1. Flow cytometry gating strategy for mouse T cells.**

(A) Debris were excluded and single cells gated.

(B) Conventional T cells (T_convs_): FoxP3^+^ T_regs_ were excluded, then CD4^+^CD8^−^ and CD8^+^CD4^−^ T cells were gated. Naïve, effector (T_eff_) and memory cells were defined as CD62L^+^CD44^-^, CD62L^-^CD44^+^ and CD62L^+^ CD44^+^, respectively.

(C) T_regs_: CD4^+^CD8^-^ T cells were gated, then CD4^+^CD8^-^CD25^+^Foxp3^+^ T_regs_ were gated.

**Figure S2. sRAGE increases the numbers of CD4+CD8-CD25+Foxp3+ Tregs and Tconvs in the pancreatic lymph nodes (PLN) and spleen on day 64.**

(A) CD4^+^CD8^-^CD25^+^Foxp3^+^ T_regs_.

(B) CD4^+^FoxP3^−^ T_convs_; and

(C) CD8^+^FoxP3^−^ T_convs_, accompanied with their CD62L^+^CD44^-^ naïve, CD62L^-^CD44^+^ effector (T_eff_) and CD62L^+^ CD44^+^ memory subsets.

Two-tailed Mann-Whitney U-test. *n* = 4-13/group. * *P* < 0.05; ** *P* < 0.01.

**Figure S3. sRAGE decreases the numbers of CD4^+^CD8^-^CD25^+^Foxp3^+^ T_regs_ and T_convs_ in the PLN and spleen on day 225.**

(A) CD4^+^CD8^-^CD25^+^Foxp3^+^ T_regs_.

(B) CD4^+^FoxP3^−^ T_convs_; and

(C) CD8^+^FoxP3^−^ T_convs_, accompanied with their CD62L^+^CD44^-^ naïve, CD62L^-^CD44^+^ effector (T_eff_) and CD62L^+^ CD44^+^ memory subsets.

Two-tailed Mann-Whitney U-test. *n* = 4-13/group. * *P* < 0.05; *** *P* < 0.001.

**Figure S4. sRAGE increases the numbers of dendritic cells and macrophages on day 64.**

(A) Conventional dendritic cells (cDCs) including CD8^+^ cDCs (CD11c^+^CD11b^-^B220^-^CD8^+^) and CD11b^+^ cDCs (CD11c^+^CD11b^+^B220^-^CD8^-^), as well as plasmacytoid dendritic cells (pDCs; CD11c^+^CD11b^-^B220^+^).

(B) Macrophages (F4/80^+^CD11c^-^CD11b^+^B220^int/hi^).

**Figure S5. Prediabetes oral glucose tolerance tests (OGTTs).**

(A-E) OGTTs on day 64 (*n* = 8/group). (F-J) OGTTs on day 80 (*n* = 15/group).

(A, F) Blood glucose concentrations; (B, G) Area under the curve for blood glucose (AUC_glucose_); (C, H) Plasma insulin concentrations; (D, I) AUC_insulin_; (E, J) AUC_glucose:insulin_ ratio.

Data shown as mean ± SD and analyzed by two-tailed unpaired Student t-tests. * *P* < 0.05 between groups.

**Figure S6. Gating strategies for human T cell proliferation experiments.**

(A and B) CD3^+^CD4^+^CD25^+^CD127^lo/–^ natural T_regs_ (nT_regs_) and CD3^+^CD4^+^ CD25^−^ T_convs_ were positively isolated by FACS.

(C and D) CD3^+^CD4^+^CD25^+^CD127^lo/–^ nT_regs_ were assessed for FoxP3 expression.

(E-G) Gating strategy for analyzing nT_reg_ and T_conv_ proliferation when co-cultured. (E) Debris, multiplets and dead cells were excluded, (F) CD3^+^CD4^+^ cells were gated. CD3^+^CD4^+^ cells were then analyzed for nT_reg_ and T_conv_ proliferation by CFSE and CellTrace Violet dilution, respectively. (G) nT_regs_ and T_convs_ were both CD25^+^CD127^lo/–^ post-stimulation, so CD25 and CD127 expression was not used to delineate T_regs_ and T_convs_ prior to CFSE and CellTrace Violet dye dilution analysis.

(H-J) Gating strategy for analyzing nT_reg_ proliferation when cultured alone. (H) Debris, multiplets and dead cells were excluded, (I) CD3^+^CD4^+^ cells were gated, (J) CD127^lo/–^ cells were gated as nT_regs_. nT_regs_ were then analyzed for proliferation by CFSE dilution.

**Figure S7. Gating strategies for human induced Treg (iTreg) differentiation experiment.**

(A and B) Purity check for naive CD4^+^ T cells (CD3^+^CD4^+^CD45RA^+^CD45RO^−^) (A) before negative isolation (peripheral blood mononuclear cells, PBMCs) and (B) after negative isolation. (C-E) Gating strategy for analyzing iT_reg_ differentiation. (C) Debris, multiplets and dead cells were excluded, (D) CD3^+^CD4^+^ cells were gated, (E) CD25^+^CD127^lo/–^ cells were gated as iT_regs_.

**Figure S8. Human iTreg differentiation in AGE-containing culture media is modestly decreased by sRAGE treatment.**

Human naive CD3^+^CD4^+^CD45RA^+^CD45RO^−^ T cells were incubated in 3 day co-culture containing 5 ng/mL IL-2, 2 ng/mL TGF-β and 1 µg/mL plate-bound anti-CD3 antibody.

(A) Representative dot plots of CD3^+^CD4^+^CD25^+^CD127^lo/–^ iT_regs_ in AGE-supplemented media with co-administration of vehicle or sRAGE (50 µg/day).

(B) Quantification of change in iT_reg_ differentiation by sRAGE treatment. Data shown as mean ± SD; paired t-tests; *n* = 4/group.

**Table S1. Human nTreg NanoString predicted upstream regulators.**

**Table S2. Human nTreg NanoString network analysis.**

**Table S3. Human nTreg NanoString volcano plot values.**

