## Supplementary Materials for "Expansion of Functional Regulatory T Cells Using Soluble RAGE Prevents Type 1 Diabetes"

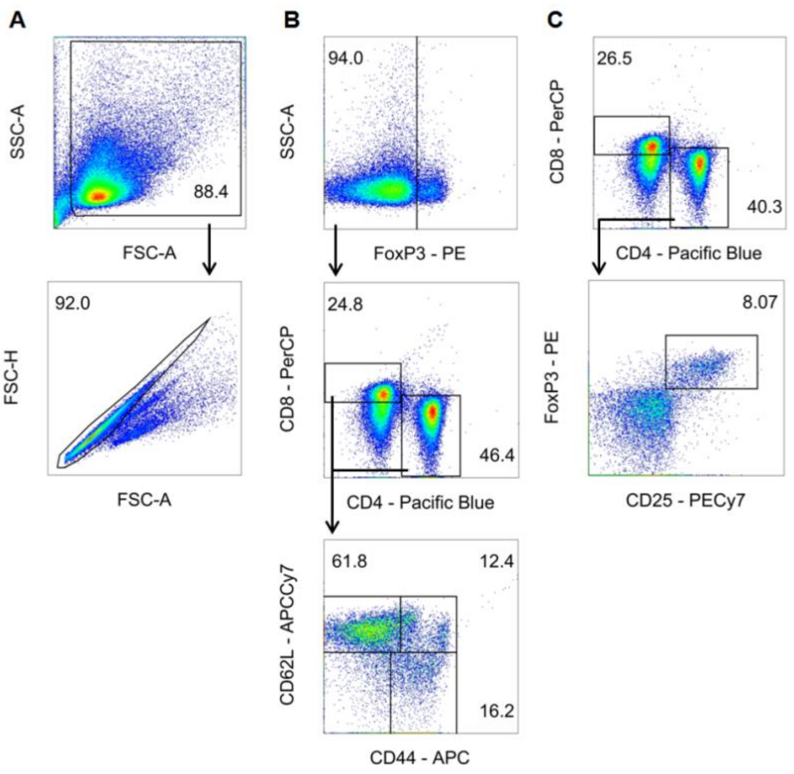


**Figure S1. Flow cytometry gating strategy for mouse T cells.**

(A) Debris were excluded and single cells gated.

(B) Conventional T cells ($T_{convs}$): FoxP3$^{+}$ $T_{regs}$ were excluded, then CD4$^{+}$CD8$^{–}$ and CD8$^{+}$CD4$^{–}$ T cells were gated. Naïve, effector ($T_{eff}$) and memory cells were defined as CD62L$^{+}$CD44$^{-}$, CD62L$^{-}$ CD44$^{+}$ and CD62L$^{+}$ CD44$^{+}$, respectively.

(C) $T_{regs}$: CD4$^{+}$CD8$^{-}$ T cells were gated, then CD4$^{+}$CD8$^{-}$CD25$^{+}$Foxp3$^{+}$ $T_{regs}$ were gated.

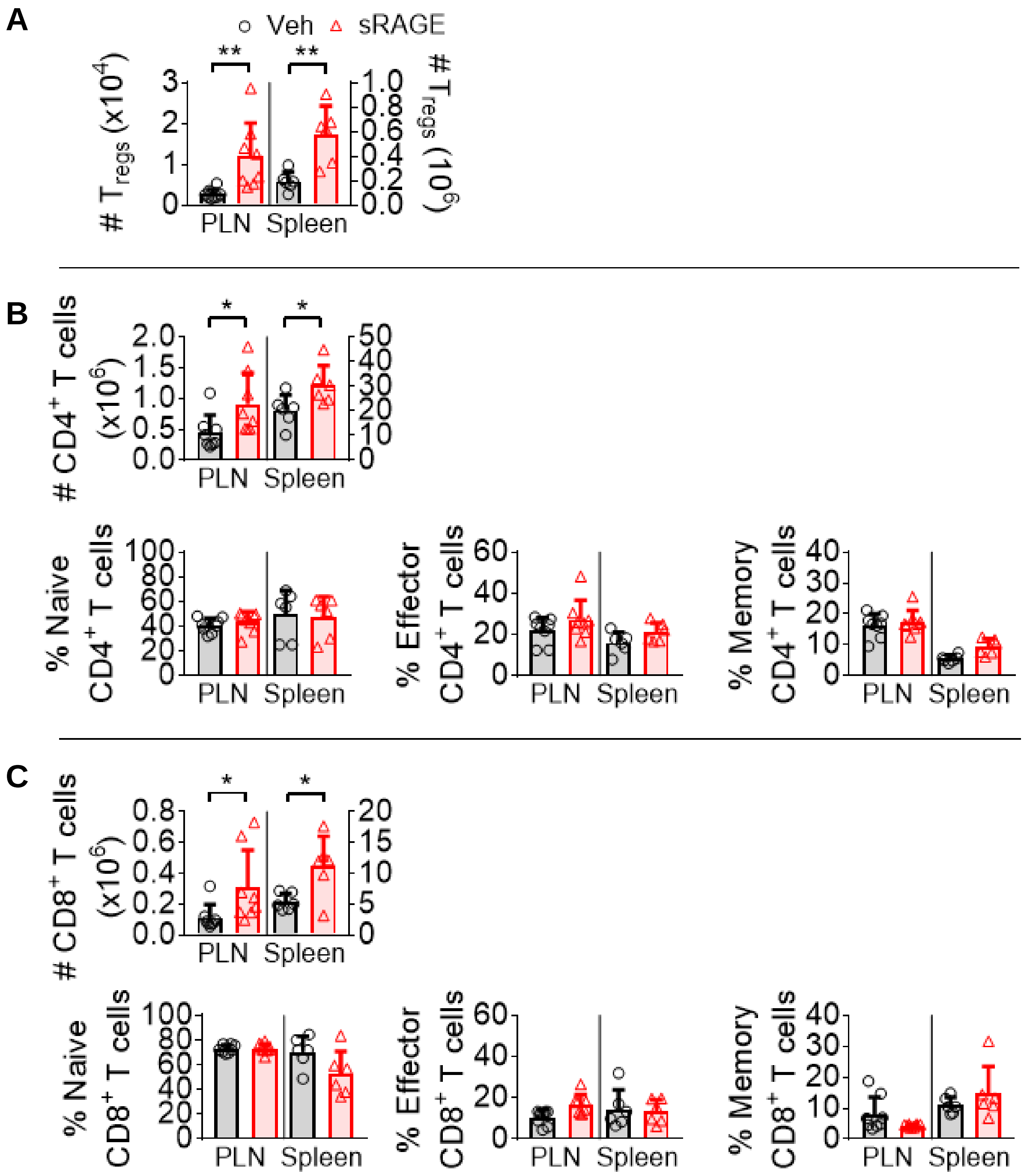


**Figure S2. sRAGE increases the numbers of CD4$^+$CD8$^-$CD25$^+$Foxp3$^+$ T$_{regs}$ and T$_{convs}$ in the pancreatic lymph nodes (PLN) and spleen on day 64.**

(A) CD4$^+$CD8$^-$CD25$^+$Foxp3$^+$ T$_{regs}$.

(B) CD4$^+$FoxP3$^-$ T$_{convs}$; and

(C) CD8$^+$FoxP3$^-$ T$_{convs}$, accompanied with their CD62L$^+$CD44$^-$ naïve, CD62L$^-$CD44$^+$ effector (T$_{eff}$) and CD62L$^+$ CD44$^+$ memory subsets.

Two-tailed Mann-Whitney U-test. *n* = 4-13/group. * *P* < 0.05; ** *P* < 0.01.

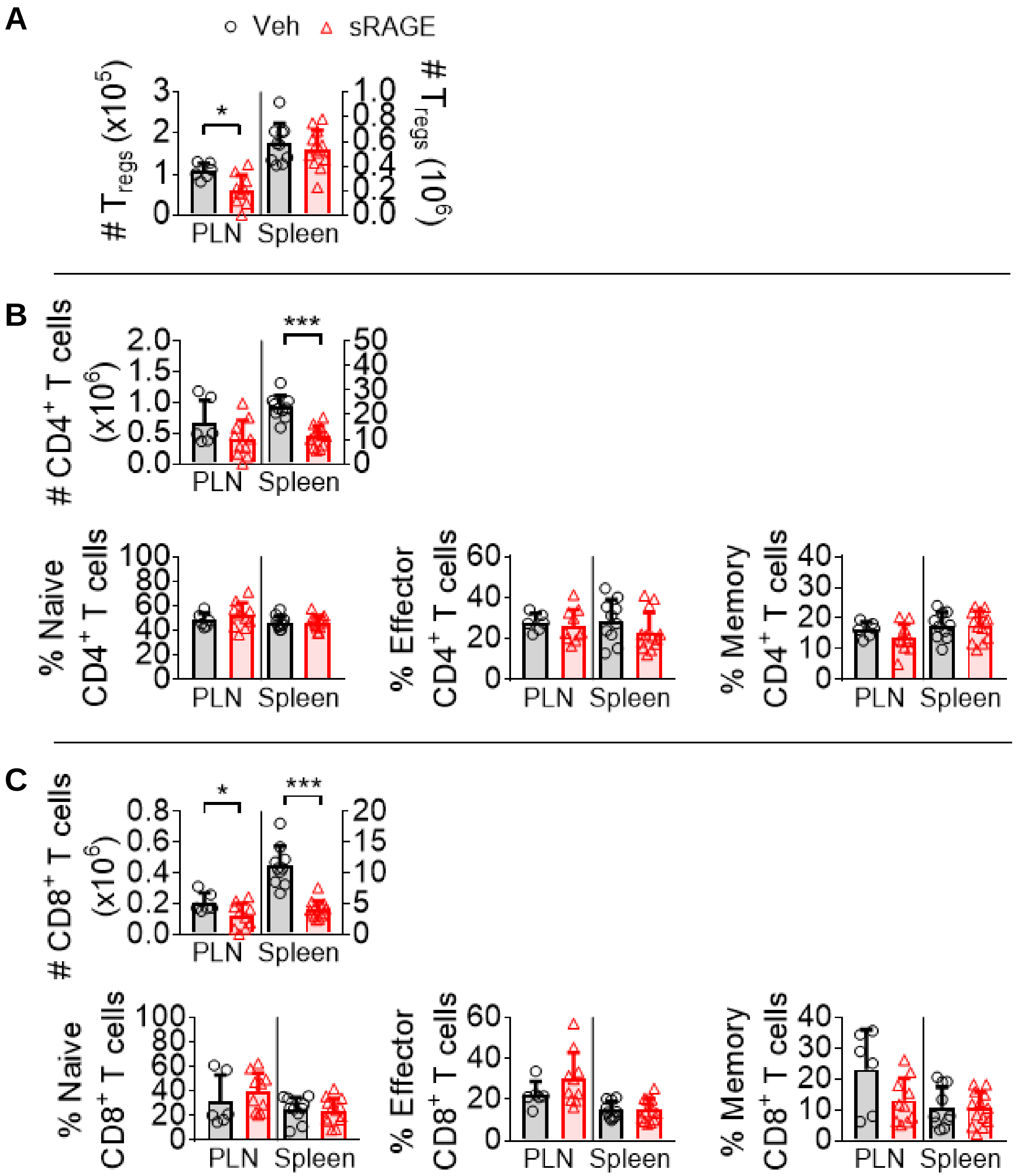


**Figure S3. sRAGE decreases the numbers of $CD4^{+}CD8^{-}CD25^{+}Foxp3^{+}$ $T_{regs}$ and $T_{convs}$ in the PLN and spleen on day 225.**

(A) $CD4^{+}CD8^{-}CD25^{+}Foxp3^{+}$ $T_{regs}$.

(B) $CD4^{+}FoxP3^{-}$ $T_{convs}$; and

(C) $CD8^{+}FoxP3^{-}$ $T_{convs}$, accompanied with their $CD62L^{+}CD44^{-}$ naïve, $CD62L^{-}CD44^{+}$ effector ($T_{eff}$) and $CD62L^{+}$ $CD44^{+}$ memory subsets.

Two-tailed Mann-Whitney U-test. $n$ = 4-13/group. * $P < 0.05$; *** $P < 0.001$.

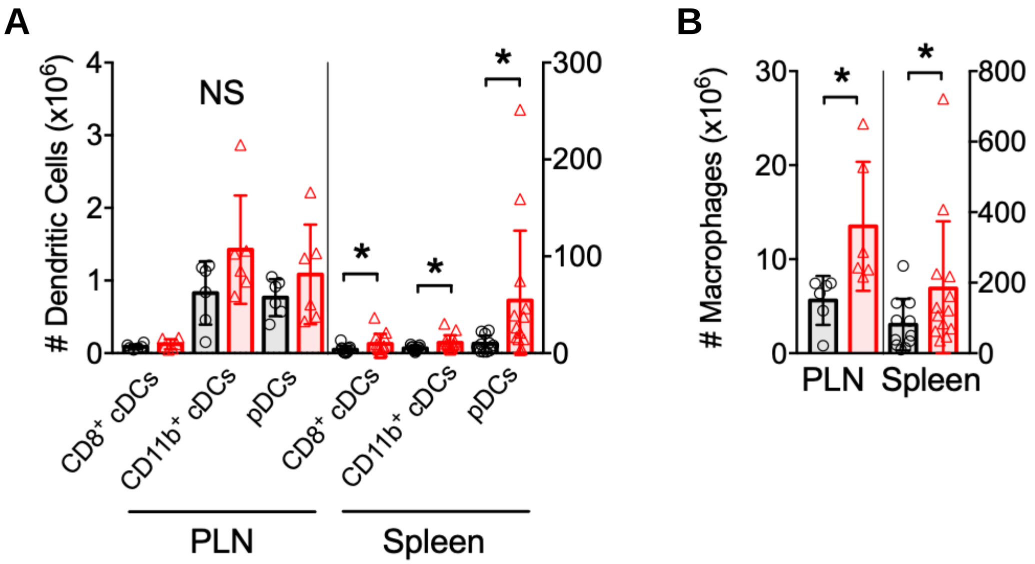


**Figure S4. sRAGE increases the numbers of dendritic cells and macrophages on day 64.**

(A) Conventional dendritic cells (cDCs) including CD8$_{+}$ cDCs (CD11c$_{+}$CD11b$_{-}$B220$_{-}$CD8$_{+}$) and CD11b$_{+}$ cDCs (CD11c$_{+}$CD11b$_{+}$B220$_{-}$CD8$_{-}$), as well as plasmacytoid dendritic cells (pDCs; CD11c$_{+}$CD11b$_{-}$B220$_{+}$).

(B) Macrophages (F4/80$_{+}$CD11c$_{-}$CD11b$_{+}$B220$_{int/hi}$).

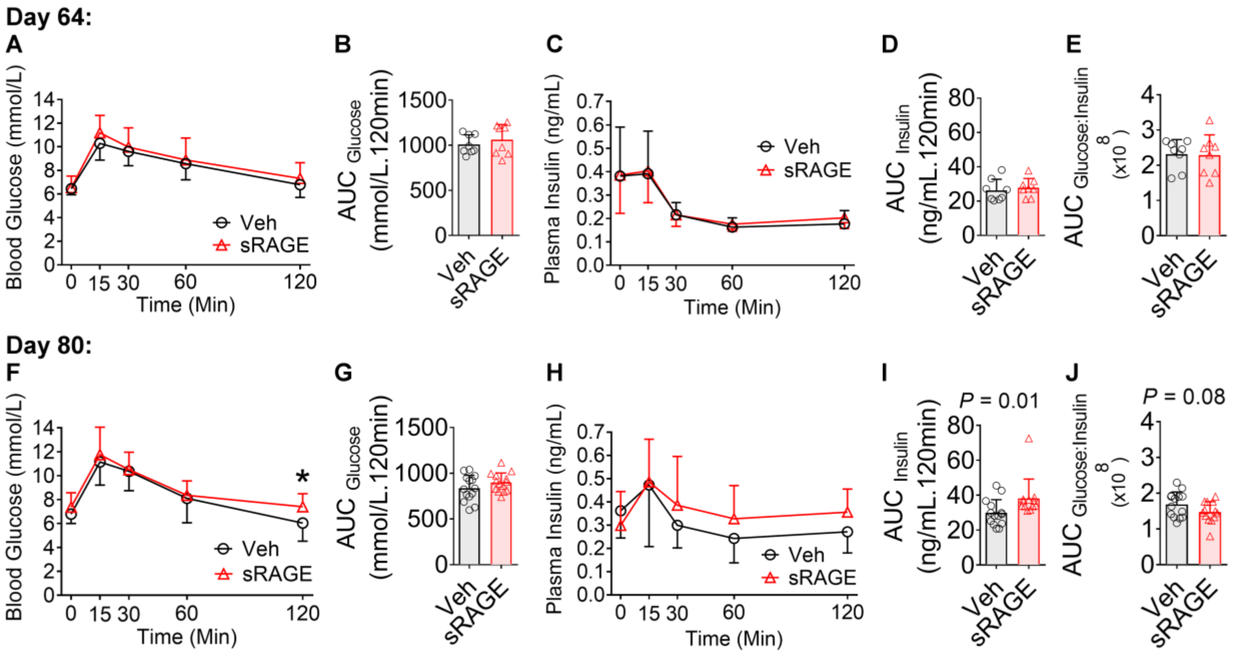


**Figure S5. Prediabetes oral glucose tolerance tests (OGTTs).**

(A-E) OGTTs on day 64 (*n* = 8/group).

(F-J) OGTTs on day 80 (*n* = 15/group).

(A, F) Blood glucose concentrations; (B, G) Area under the curve for blood glucose ($AUC_{glucose}$);

(C, H) Plasma insulin concentrations; (D, I) $AUC_{insulin}$; (E, J) $AUC_{glucose:insulin}$ ratio.

Data shown as mean ± SD and analyzed by two-tailed unpaired Student t-tests. * $P < 0.05$ between groups.

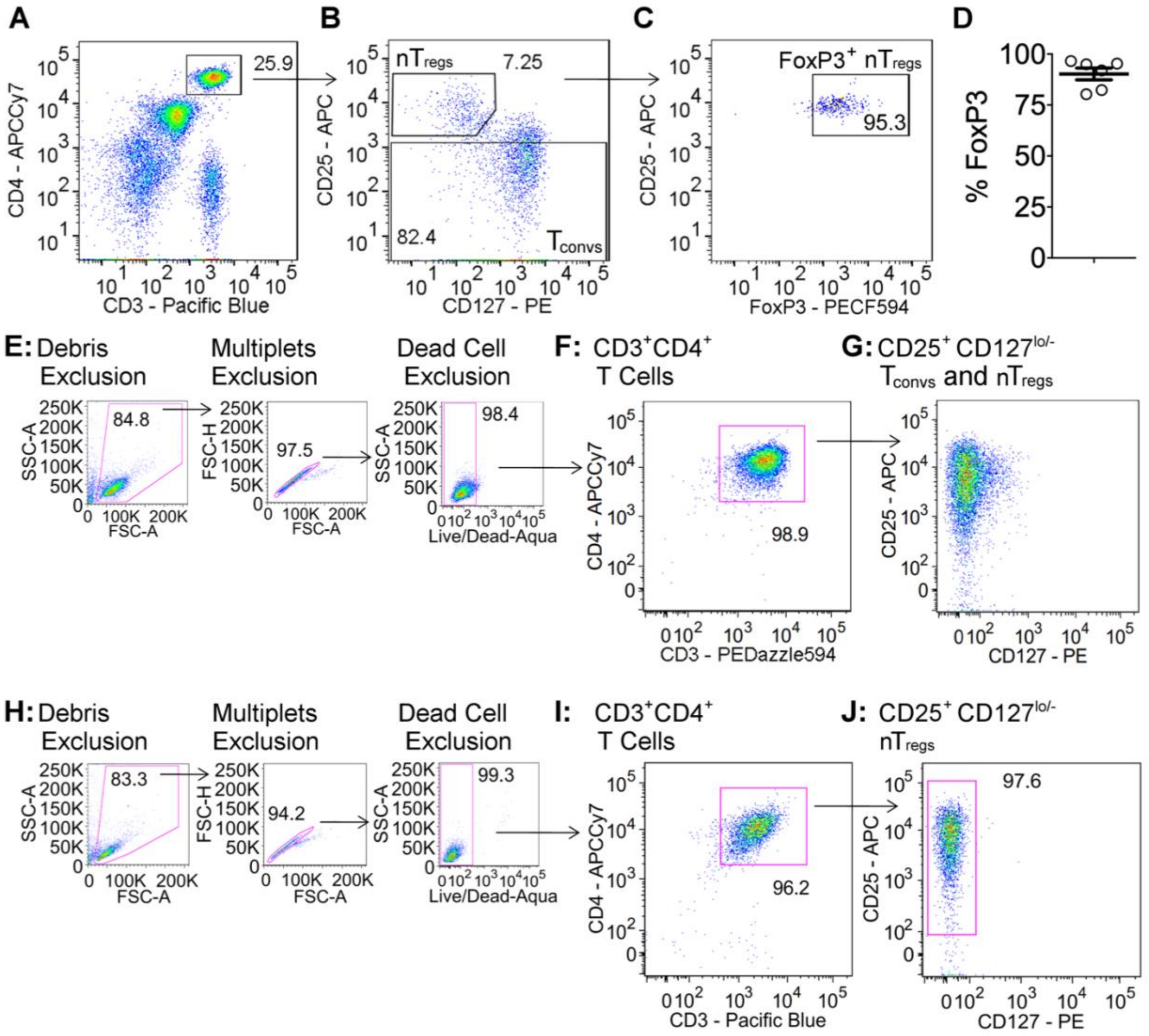


**Figure S6. Gating strategies for human T cell proliferation experiments.**

(A and B) CD3+CD4+CD25+CD127lo/– natural Tregs (nTregs) and CD3+CD4+ CD25– Tconvs were positively isolated by FACS.

(C and D) CD3+CD4+CD25+CD127lo/– nTregs were assessed for FoxP3 expression.

(E-G) Gating strategy for analyzing nTreg and Tconv proliferation when co-cultured. (E) Debris, multiplets and dead cells were excluded, (F) CD3+CD4+ cells were gated. CD3+CD4+ cells were then analyzed for nTreg and Tconv proliferation by CFSE and CellTrace Violet dilution, respectively. (G) nTregs and Tconvs were both CD25+CD127lo/– post-stimulation, so CD25 and CD127 expression was not used to delineate Tregs and Tconvs prior to CFSE and CellTrace Violet dye dilution analysis

(H-J) Gating strategy for analyzing nT$_{reg}$ proliferation when cultured alone. (H) Debris, multiplets and dead cells were excluded, (I) CD3$^{+}$CD4$^{+}$ cells were gated, (J) CD127$^{lo/–}$ cells were gated as nT$_{regs}$. nT$_{regs}$ were then analyzed for proliferation by CFSE dilution.

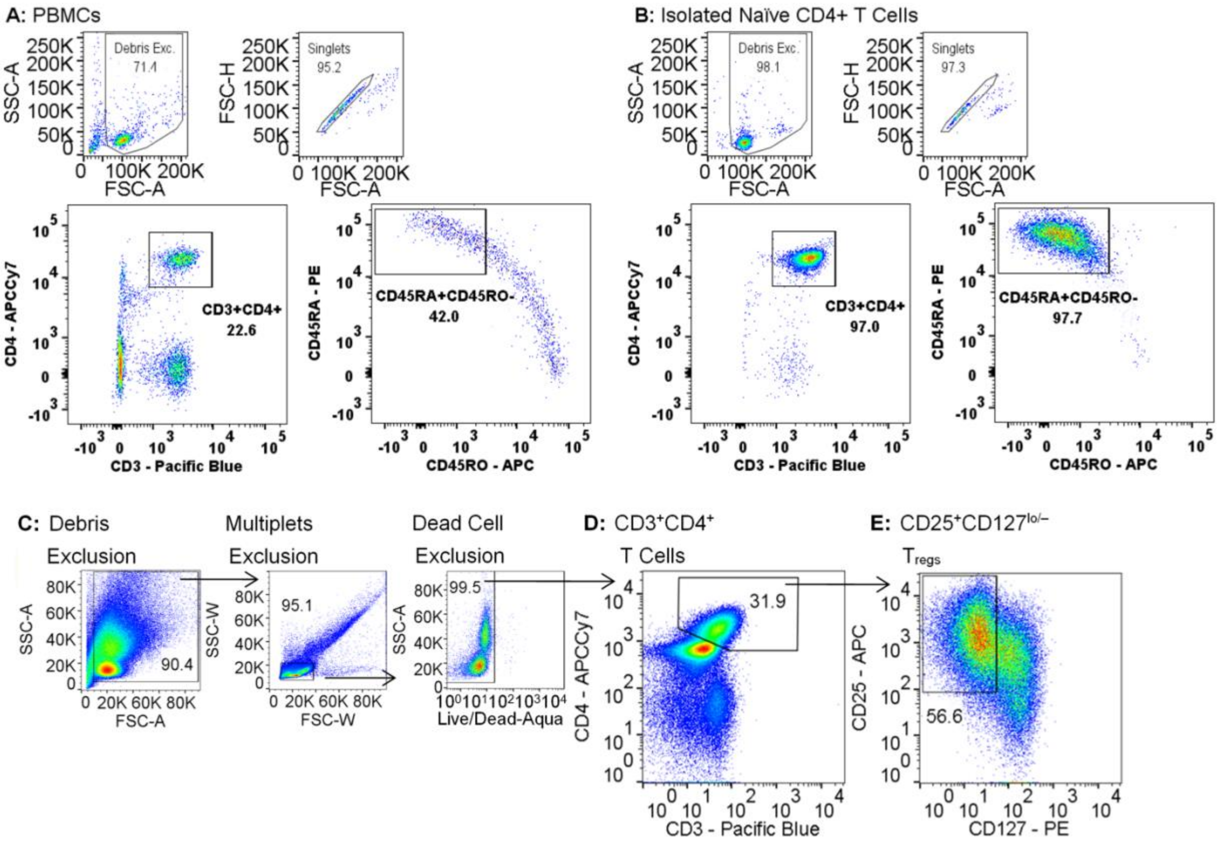


**Figure S7. Gating strategies for human induced T$_{reg}$ (iT$_{reg}$) differentiation experiment.**

(A and B) Purity check for naive CD4$^{+}$ T cells (CD3$^{+}$CD4$^{+}$CD45RA$^{+}$CD45RO$^{–}$) (A) before negative isolation (peripheral blood mononuclear cells, PBMCs) and (B) after negative isolation.

(C-E) Gating strategy for analyzing iT$_{reg}$ differentiation. (C) Debris, multiplets and dead cells were excluded, (D) CD3$^{+}$CD4$^{+}$ cells were gated, (E) CD25$^{+}$CD127$^{lo/–}$ cells were gated as iT$_{regs}$.

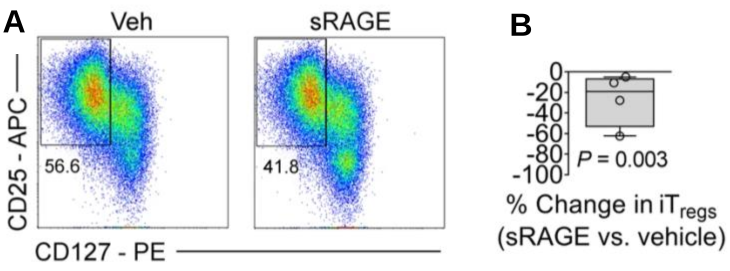


**Figure S8. Human iT$_{reg}$ differentiation in AGE-containing culture media is modestly decreased by sRAGE treatment.**

Human naive CD3$^{+}$CD4$^{+}$CD45RA$^{+}$CD45RO$^{–}$ T cells were incubated in 3 day co-culture containing 5 ng/mL IL-2, 2 ng/mL TGF-β and 1 µg/mL plate-bound anti-CD3 antibody.

(A) Representative dot plots of CD3$^{+}$CD4$^{+}$CD25$^{+}$CD127$^{lo/–}$ iT$_{regs}$ in AGE-supplemented media with co-administration of vehicle or sRAGE (50 µg/day).

(B) Quantification of change in iT$_{reg}$ differentiation by sRAGE treatment. Data shown as mean ± SD; paired t-tests; *n* = 4/group.

**Table S1. Human $nT_{reg}$ NanoString predicted upstream regulators.**

| Gene Name | Log10 FDR *P* |
|---|---|
| CD28 | 9.9136 |
| IL2 | 8.5086 |
| CD3 | 6.0650 |
| IL4 | 6.0022 |
| TBX21 | 5.5784 |
| NPC2 | 5.4168 |
| CEBPB | 4.8601 |
| STAT3 | 4.8097 |
| GATA3 | 4.5986 |
| TCR | 4.3893 |
| PDCD1 | 4.3449 |
| IL12 (complex) | 4.2197 |
| miR-9-5p (and other miRNAs w/seed CUUUGGU) | 4.2140 |
| NPC1 | 3.9830 |
| mir-9 | 3.8861 |
| TIGIT | 3.8861 |
| HSPD1 | 3.8861 |
| CHRNA3 | 3.8861 |
| TNF | 3.8297 |
| CTNNB1 | 3.7328 |
| PHB2 | 3.7190 |
| IL24 | 3.7190 |
| SAA | 3.6478 |
| TLR2 | 3.6345 |
| CD46 | 3.5186 |
| IL12B | 3.5186 |
| P2RX7 | 3.4078 |
| NR3C1 | 3.2457 |
| Interferon alpha | 3.1778 |
| FOXP3 | 3.1035 |
| NFkB (complex) | 3.1024 |
| HSF1 | 3.1002 |
| miR-21-5p (and other miRNAs w/seed AGCUUAU) | 2.9393 |
| SELPLG | 2.9393 |
| NFATC2 | 2.9393 |
| mir-155 | 2.8268 |
| POU5F1 | 2.8125 |
| REL | 2.8013 |
| VAV | 2.7620 |
| PRELID1 | 2.7620 |
| TNFRSF4 | 2.7620 |
| MIR4281 | 2.7620 |
| TPO | 2.7620 |
| TCF12 | 2.7496 |
| mir-21 | 2.7033 |
| STAT5a/b | 2.5969 |
| TGFB1 | 2.5376 |
| RELA | 2.4724 |
| BTK | 2.4685 |
| PIWIL2 | 2.4597 |
| USP21 | 2.4597 |
| IL21 | 2.2993 |
| TLR9 | 2.2993 |
| IL10RA | 2.2840 |
| FLOT2 | 2.2840 |
| ICOSLG/ LOC102723996 | 2.2840 |
| PTPN22 | 2.2840 |
| BCR (complex) | 2.2588 |
| COL18A1 | 2.2328 |
| HIF1A | 2.2218 |
| calpain | 2.1599 |
| CRBN | 2.1599 |
| MAP2K4 | 2.1599 |
| COPA | 2.1599 |
| IL4R | 2.1599 |
| IL10 | 2.0926 |
| ELAVL1 | 2.0814 |
| Integrin | 2.0630 |
| HIF3A | 2.0630 |
| PIM3 | 2.0630 |
| ZBTB7B | 2.0630 |
| SPIB | 2.0630 |
| BACH2 | 2.0630 |
| MYC | 2.0462 |
| MTOR | 2.0205 |
| CYTOR | 1.9830 |
| TNFRSF8 | 1.9830 |
| CTBP2 | 1.9830 |
| ALK | 1.9830 |
| NANOG | 1.9469 |
| ETS | 1.9172 |
| PRKDC | 1.9172 |
| NR4A2 | 1.9172 |
| STAT5B | 1.9172 |
| WNT3A | 1.9172 |
| CAMK4 | 1.8601 |
| TNFSF9 | 1.8601 |
| SSTR2 | 1.8601 |
| CDKN1B | 1.8601 |
| EGFR | 1.8210 |
| CLEC7A | 1.8097 |
| TPM3 | 1.8097 |
| mir-132 | 1.8097 |
| STK4 | 1.8097 |
| IL9 | 1.8097 |
| PTPN6 | 1.8097 |
| RPS6KA3 | 1.8097 |
| CYP1B1 | 1.8097 |
| BRD4 | 1.7696 |
| Wnt | 1.7645 |
| BCL6B | 1.7645 |
| CD5 | 1.7645 |
| FCGR2A | 1.7235 |
| PIM2 | 1.7235 |
| ENG | 1.7235 |
| IL7 | 1.7235 |
| ELF5 | 1.7235 |
| CEBPA | 1.6968 |
| mir-154 | 1.6861 |
| NR6A1 | 1.6861 |
| CFLAR | 1.6861 |

| | |
|---|---|
| PRKCD | 1.6840 |
| IFNG | 1.6737 |
| IL6 | 1.6655 |
| CRTC2 | 1.6517 |
| IL12 (family) | 1.6517 |
| LRP6 | 1.6517 |
| SYN1 | 1.6517 |
| ICOS | 1.6517 |
| TMSB4 | 1.6198 |
| Calcineurin protein(s) | 1.6198 |
| NFKBIZ | 1.6198 |
| RORC | 1.6198 |
| MIF | 1.6198 |
| TRIB3 | 1.5901 |
| FOXP1 | 1.5901 |
| Tgf beta | 1.5622 |
| DPH5 | 1.5622 |
| PRDM1 | 1.5622 |
| BCL11B | 1.5622 |
| USP7 | 1.5361 |
| CAV1 | 1.5361 |
| RAC1 | 1.5114 |
| MMP2 | 1.5114 |
| TP73 | 1.4935 |
| Smad2/3 | 1.4881 |
| TNFRSF1A | 1.4881 |
| KIT | 1.4881 |
| Mapk | 1.4461 |
| GFI1 | 1.4461 |
| ZBTB16 | 1.4260 |
| RAF1 | 1.4260 |
| ATF2 | 1.4260 |
| CCL2 | 1.4067 |
| ZEB1 | 1.4067 |
| NR1I2 | 1.4067 |
| USF2 | 1.4067 |
| PIM1 | 1.3893 |
| CTLA4 | 1.3716 |
| RELB | 1.3716 |

| | |
|---|---|
| SMARCA4 | 1.3706 |
| Hsp27 | 1.3546 |
| F7 | 1.3546 |
| NFKB2 | 1.3546 |
| RNASE2 | 1.3391 |
| RARA | 1.3363 |
| TAL1 | 1.3279 |
| miR-155-5p (miRNAs w/ seed UAAUGCU) | 1.3233 |
| mir-29 | 1.3233 |
| HMGA1 | 1.3089 |
| PKM | 1.3089 |

**Table S2. Human $nT_{reg}$ NanoString network analysis.**

| Focus Molecules (Count / Total) | Log10 *P* (Score) |
|---|---|
| AHR, GZMA, JAK1, STAT4, BACH2, FOXP3, GZMB, IL23A, JAK3, TIGIT, CCR4, GATA3, IL10RA, IL4R, IRF4, SOCS5, STAT5B, TNFRSF1A, CD226, IL7R, ITK, NFATC2, STAT3, STAT6 (24/44 molecules) | 57 |
| BTLA, KLF3, LGALS1, TNF, CCR4, CCR7, EOMES, GZMK, PTGER2 (9/35 molecules) | 11 |
| CD96 (1/2 molecules) | 2 |
| GNLY, PDCD1 (2/5 molecules) | 2 |
| IFNG, KLRB1, ZBTB16 (3/8 molecules) | 2 |
| CD27, CD40LG (2/12 molecules) –*not significant* | 1 |
| ITGA1, TGFB1 (2/22 molecules) – *not significant* | 1 |

**Table S3. Human $nT_{reg}$ NanoString volcano plot values.**

| Gene Name | Log2 fold change | Log10 FDR *P* |
|---|---|---|
| **Downregulated Genes** | | |
| ***Significant & Log2 Fold Change > \|1\|*** | | |
| IL7R | -2.5484 | -1.7546 |
| CD49a | -2.0803 | -2.0649 |
| PTGER2 | -1.9939 | -2.1195 |
| TIGIT | -1.8914 | -2.9172 |
| IL23A | -1.5512 | -2.0649 |
| BACH2 | -1.4216 | -1.3429 |
| CD96 | -1.3483 | -2.2805 |
| IL10RA | -1.2758 | -1.8328 |
| GZMK | -1.2721 | -1.5859 |
| EOMES | -1.1603 | -2.2805 |
| STAT5B | -1.0845 | -2.2805 |
| STAT4 | -1.0534 | -2.2805 |
| SOCS5 | -1.0118 | -2.2805 |
| AHR | -1.0036 | -2.2805 |
| KLRB1 | -1.0012 | -1.6611 |
| ***Significant & Log2 Fold Change < \|1\|*** | | |
| JAK1 | -0.9607 | -2.2805 |
| ITK | -0.9462 | -2.2399 |
| CD27 | -0.9205 | -1.5859 |
| GNLY | -0.9122 | -1.7023 |
| CD272 | -0.8973 | -1.7546 |
| GATA3 | -0.8789 | -2.2805 |
| IRF4 | -0.8451 | -1.7023 |
| CCR4 | -0.7954 | -2.2274 |
| STAT6 | -0.7749 | -2.0649 |
| KLF3 | -0.7526 | -1.3783 |
| CD226 | -0.7137 | -1.9163 |
| FOXP3 | -0.6978 | -1.5387 |
| TNFR1 | -0.6097 | -1.4873 |
| STAT3 | -0.6097 | -1.3095 |
| NFATC2 | -0.6054 | -1.7023 |
| IL4R | -0.5634 | -1.4870 |
| JAK3 | -0.5127 | -1.3019 |
| ***Not Significant*** | | |
| EGR2 | -1.8547 | -0.9349 |
| BCL6 | -1.6249 | -1.1240 |
| IL9 | -1.6055 | -0.2823 |
| NT5E | -1.3630 | -1.2081 |
| ENTPD1 | -1.2501 | -0.8104 |
| IL1A | -1.1942 | -1.2454 |
| IFNAR1 | -0.9949 | -1.1934 |
| JUN | -0.9625 | -0.8104 |
| CD137 | -0.9331 | -1.0007 |
| CD45RA | -0.9205 | -0.2759 |
| CD134 | -0.8898 | -0.8104 |
| CXCR5 | -0.8268 | -0.6385 |
| CCR6 | -0.7857 | -1.1781 |
| ADORA2A | -0.7784 | -1.2081 |
| CD69 | -0.7732 | -1.0007 |
| FOS | -0.7692 | -0.4594 |
| CCL5 | -0.7610 | -0.5731 |
| IL18R1 | -0.7587 | -0.9195 |
| CXCR6 | -0.7412 | -1.1861 |
| MAF | -0.7076 | -0.8492 |
| CCR7 | -0.6972 | -0.9349 |
| JAK2 | -0.6870 | -1.1781 |
| ITGA4 | -0.6773 | -0.8492 |
| CXCL8 | -0.6668 | -0.2232 |
| CX3CR1 | -0.6629 | -0.6085 |
| KLF2 | -0.6381 | -1.0614 |
| IL21R | -0.6134 | -0.5089 |
| HLA.DRA | -0.5774 | -1.2081 |
| IL27RA | -0.5764 | -1.1781 |
| PRDM1 | -0.5591 | -0.8104 |
| IFNGR1 | -0.5530 | -1.2081 |
| STAT5A | -0.5115 | -1.0310 |
| GZMM | -0.5023 | -0.1776 |
| IL2RB | -0.5011 | -0.9057 |
| SOCS1 | -0.4920 | -1.0009 |
| PTGDR2 | -0.4850 | -0.5087 |
| ZBTB16 | -0.4826 | -0.4766 |
| STAT1 | -0.4627 | -1.1781 |
| CD45R0 | -0.4396 | -0.8966 |
| SELL | -0.4279 | -1.1240 |

| | | |
|---|---|---|
| RUNX3 | -0.4201 | -0.9349 |
| RUNX1 | -0.4166 | -1.2454 |
| IL32 | -0.3852 | -1.2569 |
| PVRIG | -0.3722 | -0.6077 |
| CTLA4 | -0.3668 | -0.5973 |
| IL13 | -0.3609 | -0.1390 |
| CCR5 | -0.3567 | -0.1867 |
| IL12RB2 | -0.3445 | -0.7985 |
| CCL4 | -0.3087 | -0.1074 |
| CD3D | -0.3003 | -1.0009 |
| FAS | -0.2993 | -0.7200 |
| NFKB1 | -0.2641 | -0.3327 |
| CD4 | -0.2562 | -0.8104 |
| GFI1 | -0.2466 | -0.5371 |
| PDCD1 | -0.2256 | -0.2111 |
| IL2RA | -0.1373 | -0.2670 |
| TGFB1 | -0.1181 | -0.2759 |
| LCK | -0.1157 | -0.2823 |
| CD28 | -0.0971 | -0.2681 |
| IFNG | -0.0852 | -0.0887 |
| TNF | -0.0831 | -0.0751 |
| S1PR1 | -0.0366 | -0.0942 |
| PRF1 | -0.0214 | -0.0208 |
| SOCS3 | 0.0052 | -0.0034 |
| **Upregulated Genes** | | |
| ***Significant & Log2 Fold Change > \|1\|*** | | |
| GZMB | 2.4512 | -2.1857 |
| GZMA | 1.8410 | -2.2805 |
| LGALS1 | 1.0369 | -2.1817 |
| ***Not Significant*** | | |
| IL10 | 1.6348 | -1.2429 |
| IL17F | 0.9889 | -0.5126 |
| IL4 | 0.9328 | -0.9304 |
| CXCR3 | 0.8543 | -1.0007 |
| IL22 | 0.8312 | -0.4766 |
| HAVCR2 | 0.4377 | -1.1479 |
| LAG3 | 0.3871 | -0.5112 |
| CD40LG | 0.3713 | -0.8358 |
| ITGAE | 0.2601 | -0.6085 |
| TBX21 | 0.2408 | -0.4799 |
| CD38 | 0.2017 | -0.2113 |
| ICOS | 0.1964 | -0.4859 |
| RORC | 0.1478 | -0.2113 |
| IL5 | 0.1379 | -0.0361 |
